# Genomic and pan-genomic insight into longan domestication and improvement

**DOI:** 10.64898/2026.01.04.697543

**Authors:** Jingxuan Wang, Jing Wang, Jie Sun, Jianguang Li, Ke Wang, Shaoying Chen, Xiangfeng Wang, Xiaoliang Ren, Junbin Wei, Jingyan Li, Li Guo

**Affiliations:** Peking University Institute of Advanced Agricultural Sciences, Shandong Laboratory of Advanced Agricultural Sciences in Weifang, Shandong 261325, China; Institute of Fruit Tree Research, Guangdong Academy of Agricultural Sciences; Key Laboratory of South Subtropical Fruit Biology and Genetic Resource Utilization, Ministry of Agriculture and Rural Affairs; Guangdong Provincial Key Laboratory of Science and Technology Research on Fruit Trees, Guangzhou, 510640 China; State Key Laboratory of Wheat Improvement, College of Life Sciences, Shandong Agricultural University, Tai’an, Shandong 271018, China

## Abstract

Longan (*Dimocarpus longan*) is a tropical tree in Sapindaceae family with economic importance known for its nutritous fruits. Genetic improvement of longan requires knowledge of reliable molecular markers and functional genes associated with key traits. However, it is largely impeded by lacking a complete genome and pangenome reference encompassing the broad genetic diversity in diverse longan germplasms. Here, we present a telomere-to-telomere (T2T) gap-free genome of longan cultivar ’Shixia’ and a graph-based pangenome constructed using newly assembled chromosome-level genomes of 101 accessions. We completely assemble the longan centromere regions primarily composed of Gypsy-CRM retrotransposons. The pan-genome analysis reveals 58,978 non-redundant structural variants (SVs) that exhibit signs of genomic selective sweep during longan domestication and breeding. Additionally, haplotype-resolved genome analysis suggests allele-specific gene expression associated with SV-driven changes in 3D genome architecture. Importantly, the graph-based pan-genome empowers population-scale SV genotyping and genomewide association with various longan traits. SV-GWAS revealed 12 QTLs significantly linked with longan maturity period. Among them, a 758bp insertion is located downstream of the *DlDAZ* gene encoding a C2H2 zinc-finger transcription factor, and *DIDAZ-*transgenic tomatoes showed delayed maturity. Together, our T2T genome and pan-genome provide valuable resources to facilitate longan genetic research and precise improvement.

## Introduction

Longan (*Dimocarpus longan* Lour.), an evergreen fruit tree species within the Sapindaceae family that includes the lychee and yellowhorn, is an economically vital crop extensively cultivated in tropical and subtropical regions. Domesticated from its wild relatives around 2000 years ago in China^1^, cultivated longan has been utilized in both traditional and modern medicine. It was honored as the "king of fruits" by Li Shizhen, a renowned expert in Traditional Chinese Medicine from the Ming Dynasty. Current global production of longan shows significant geographical concentration, with three Asian nations China, Thailand, and Vietnam leading commercial cultivation. China leads in both cultivation area and yield^2^, while longan was introduced to the United States in late 19th century. The maturity period of fruit is a key factor affecting its quality and market value^3^. Fruits that mature very early or very late typically have higher market prices and yields greater economic benefits. Therefore, understanding the mechanisms underlying fruit development, especially the maturity period, has been a key goal of research. However, such studies in longan remain limited, and the potential of longan germplasm for maturity period breeding has yet to be fully exploited due to the lack of a complete longan genome.

The initial draft genome of longan was assembled for the "Honghezi" cultivar in 2017^4^, followed by a chromosome-level genome assembly of the "Jidanben" cultivar in 2022^5^. However, numerous gaps existed and the centromeres and telomeres regions were still unresolved. Moreover, the genomic resources available for longan germplasm are insufficient, limiting the precise identification of genomic variants and the comprehensive resolution of centromeres and telomeres. Several studies have highlighted that reliance on a single reference genome significantly impacts the effectiveness of identifying intraspecific genetic variations. This is primarily because reference mapping failed to capture novel or highly divergent sequences within a species^6^. SVs including large deletions, insertions, duplications, and chromosomal rearrangements, constitute major sources of genetic diversity^7,8^. Increasing evidence demonstrates the pivotal role of SVs in plant evolution and agriculture, influencing traits such as panicle architecture, flowering time, fruit quality, and stress resistance^8–11^. To overcome these limitations, pan-genome analyses based on multiple *de novo* assembled genomes have been widely adopted to capture a broader spectrum of genetic diversity, and enrich understanding of genome evolution and plant breeding^12^.

In this study, we generate a T2T genome assembly for the longan cultivar ’Shixia’, one of the predominant commercially cultivated longan varieties renowned for its early mature period and superior fruit quality in China. We then integrate newly assembled high-quality chromosome-level genomes from 100 other longan accessions to construct the most comprehensive longan pan-genome to date. We assessed genome-wide SV diversity in all 101 longan accessions and investigated genomic evolution through domestication and breeding of longan. Pan-3D genomic analysis revealed that both SVs and 3D genomic structure affect allele-specific expression (ASE) of genes. Furthermore, we conducted SNP- and SV-GWAS to investigate agronomic traits related to fruit quality and maturity period, revealing 12 significant SV-QTLs, one of which affects the *DlDAZ* gene which encodes C2H2 zinc-finger transcription factor, and *DIDAZ* transgenic tomatoes showed delayed maturity. Together, the longan T2T genome and pan-genome provide valuable resources for accelerating its genetic research and precision improvement.

## Results

### T2T gap-free assembly of longan ’ShiXia’ genome

We first assembled a T2T genome of a major longan (*D. longan*) cultivar ’Shixia’ as a reference (**Fig. 1a**). We generated a combination of 112.55Gb (∼258x) PacBio high-fidelity (HiFi) reads, 34.17Gb (∼78x) Oxford Nanopore Technologies ultra-long (ONT-UL) reads, and 56.09 Gb high-throughput chromatin conformation capture (Hi-C) reads for ’Shixia’ (**Table S1**). The draft genome was assembled using both HiFi and ONT-UL reads, achieving a contig N50 of 28.34Mb with hifiasm^13^. The draft genome was further anchored onto 15 chromosomes using Hi-C reads with Juicer^14^ and 3d-DNA pipeline^15^, followed by manual corrections in Juicebox^16^. Following the gap-filling process with ONT reads, the final ’Shixia’-T2T assembly was 436Mb comparable to the predicted genome size using k-mer frequency (**Fig. S1a**). We captured all 15 centromeres and most (28/30) telomeres, and filled all 1189 gaps present in the previous ’Shixia’ genome assembly (117.71 kb; **Fig. 1b, Fig. S2a-b**), representing the most complete longan genome so far.

**Figure 1.**
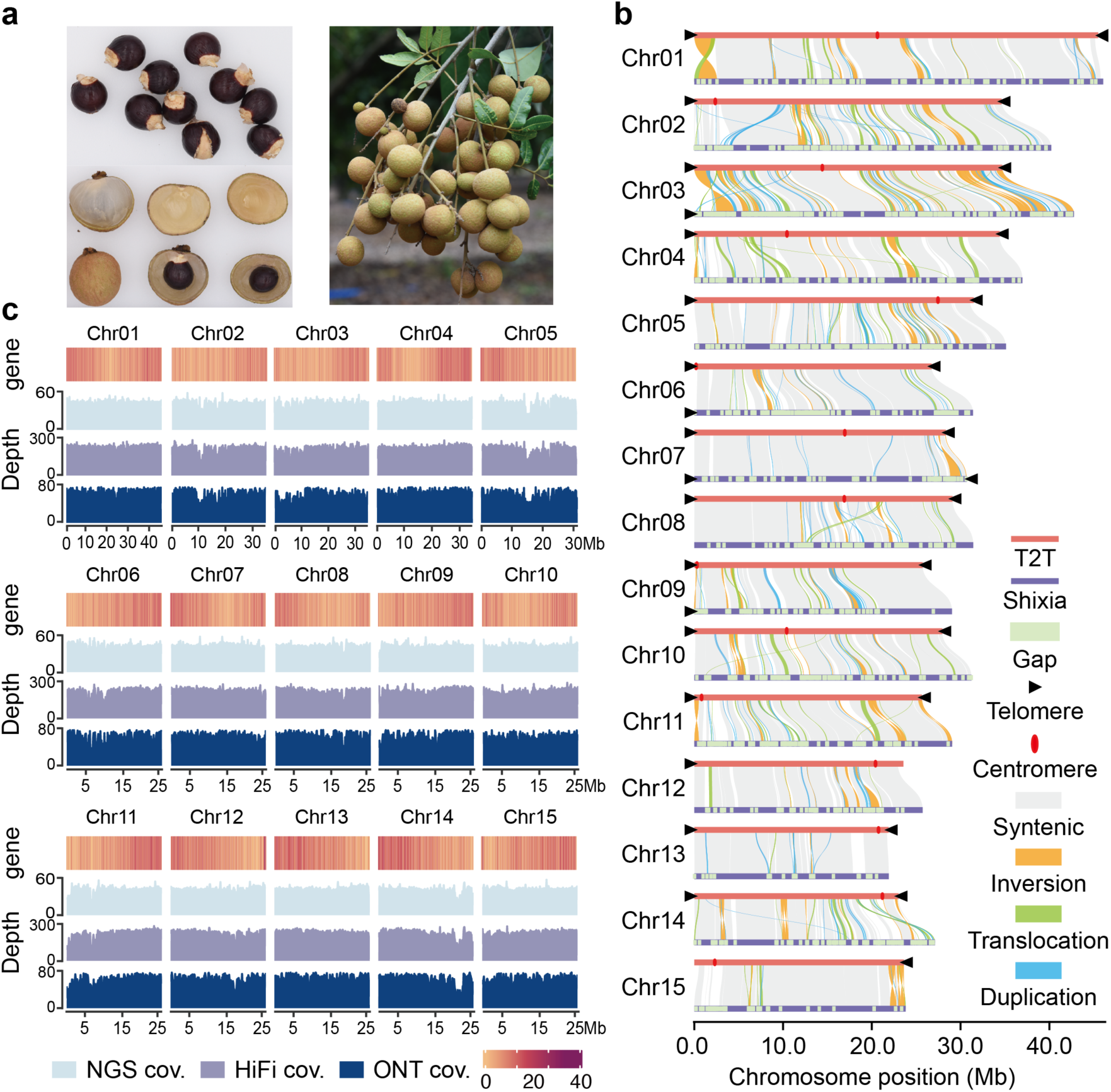
A T2T gap-free reference genome of longan cultivar ’Shixia’. **a**, A photograph of ’Shixia’ fruit section and fruit cluster. **b**, Genomic alignment between the T2T genome and the previous version of ’Shixia’ genome (ASM2045787v1) shows synteny (grey lines) and structural variants. The light green rectangle indicates the position of gaps, the red dot indicates the centromere, the black triangle marks the telomere, the green line represents translocation, and the yellow line represents inversion, all labeled at their corresponding positions. **c**, Whole-genome gene density and coverage of mapped sequencing reads generated by different sequencing technologies (NGS, HiFi and ONT) across each chromosome.

Next, we extensively validated the ’Shixia’ T2T genome using multiple methods. First, Hi-C read mapping and contact maps showed no obvious structural errors in the ’Shixia’ T2T assembly (**Fig. S1b**). Second, the mapping rates of HiFi and ONT reads to the ’Shixia’ T2T assembly were 99.28% and 99.89%, respectively, both showing overall uniform read mapping coverage across the genome (**Fig. 1c**). Finally, the ’Shixia’ T2T genome assembly displayed high completeness and base-level accuracy, with a Benchmark Universal Single Copy Ortholog (BUSCO) score of 99.70%, a LTR Assembly Index (LAI) of 15.90, and a Quality Value (QV) of 70.29. In addition, a total of 36083 protein-coding genes were annotated for the ’Shixia’ T2T assembly by integrating *ab initio* predictions, transcriptome sequencing data (**Table S2**), and homology evidence, resulting in improved gene structure accuracy compared to previous reports (**Fig. S1c**). Overall, this ’Shixia’ genome is the first T2T gap-free longan genome which will serve as a valuable resource for accelerating longan genetics and breeding.

### Characterization of centromere architecture in longan cultivar ’Shixia’

Centromeres are specialized chromosomal domains that attach to spindle microtubules, ensuring faithful chromatid segregation in cell division. We determined the centromeric regions of the longan genome based on CENH3 chromatin immunoprecipitation followed by sequencing (ChIP-seq) (**Fig. 2b**). The average size of centromeres was 400 kb, ranging from 160 kb (Chr02) to 962 kb (Chr01) (**Fig. 2b; Table S5**). Karyotypic analysis based on q/p arm ratios revealed that longan chromosomes predominantly exhibit telocentric and metacentric configurations (10m+2sm+2st+16t) (**Fig. S3a**). Repeat composition analysis demonstrated that longan centromeres were primarily composed of LTR retrotransposons (76.67%), of which *Gypsy* constituted the predominant fraction (73.39%) with minor DNA transposon components (8.44%) (**Fig. 2a**). Centromeric regions showed elevated GC content compared to non-centromeric regions. Notably, unlike the satellite-repeat dominated centromeres in humans and ape genomes^17^, longan centromeres exhibited significantly reduced non-B DNA sequence proportions (**Fig. 2a**). In addition to repeat sequences, we annotated 40 genes in longan centromeres, with functional enrichment in biological processes such as protein folding, heat stress response, and fatty acid metabolism, particularly through chaperone-mediated pathways and ATPase regulatory activities (**Fig. S3a, Fig. S4**). LTR subfamily classification identified *CRM* (*Gypsy* subfamily) as the most abundant subtype (53.85%) in centromeres. Phylogenetic analysis using reverse transcriptase (RT) domains from intact *CRM* elements revealed widespread recent integration events genome wide. Interestingly, phylogeny of longan *CRM* elements lacked apparent differentiation between centromeric and non-centromeric regions (**Fig. S3b**), indicating a non-preferential proliferation. However, centromeres exhibited significant enrichment of intact LTR sequences, predominantly occupied by *CRM* elements (**Fig. 2c**), suggesting slower turnover of *CRM* subtype LTRs within centromeres, likely under epigenetic regulation. Consistently, LTR integrity truncation ratios demonstrated a significantly higher structural preservation of *CRM* elements in centromeric regions (**Fig. 2c**). Finally, while CENH3 ChIP-seq detected comparable CENH3 recruitment for truncated *CRM* and other truncated LTRs, intact *CRM*s showed significantly stronger CENH3 enrichment than truncated *CRMs* (**Fig. 2d, Fig. S3c**). Collectively, the predominance and structural integrity of *CRM* retrotransposons in longan centromeres was likely essential for its centromere function maintenance.

**Figure 2.**
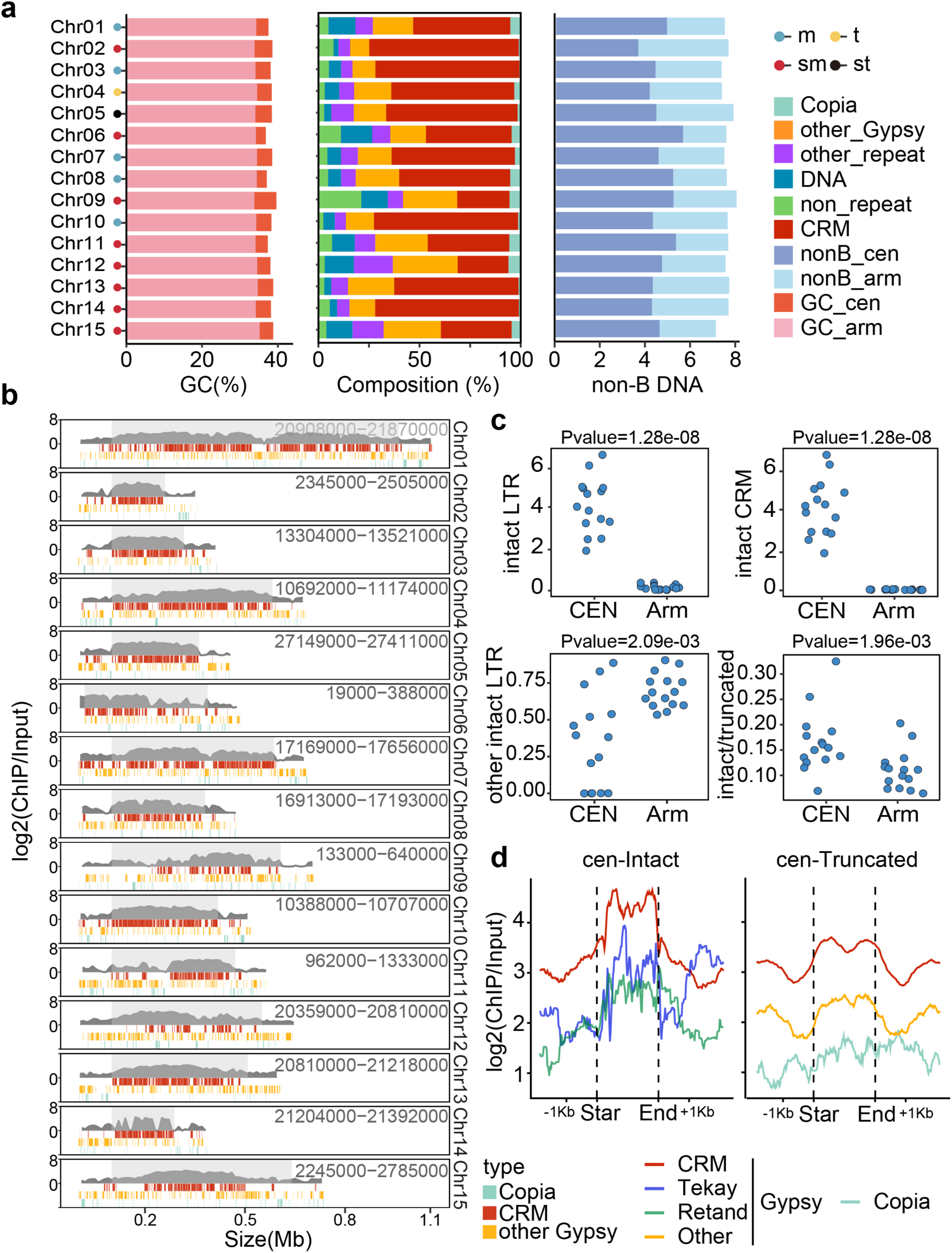
Landscape of the longan centromere repeats dominated by Gypsy-CRM retrotransposons. **a**, GC content, TE composition, and non-B DNA density (per kilobase) for each chromosome. **b**, Depiction of each centromere and flanking 100 kb regions. The shaded grey box represents the centromeric core. Shown within are: the CENH3 ChIP-seq enrichment profile (stacked peaks), and the distribution of major LTRs. **c**, Comparative analysis of LTR distribution in centromeric versus non-centromeric regions(subpanels left-right, top-bottom); (i) Density distribution of intact LTRs (per 100 kb) in centromeric vs. non-centromeric regions; (ii) Density distribution of intact CRMs (per 100 kb) in centromeric vs. non-centromeric regions; (iii) Density distribution of LTRs excluding CRM (per 100 kb) in centromeric vs. non-centromeric regions; (iv) Comparative Integrity/Truncation (I/T) ratio of CRMs in centromeric vs. non-centromeric regions. **d**, Comparative ChIP-seq enrichment of repeat elements in centromeric versus non-centromeric regions (Subpanels arranged left-to-right and top-to-bottom); (i) Genome-wide ChIP enrichment profiles of intact LTRs; (ii) Genome-wide ChIP enrichment profiles of truncated LTRs.

### Pangenome analysis from chromosome-level genomes of 101 longan accessions

To establish a comprehensive longan pangenome encompassing a wide range of genetic diversity, we collected 100 additional genetically diverse longan accessions (**Fig. 3a**) with remarkable phenotypic diversity (**Fig. 3b**). We then sequenced them using PacBio and ONT long-read technologies, where nine elite accessions were sequenced using PacBio HiFi, and 91 other accessions for 28x ONT and 67x NGS sequencing (**Table S1**), and then performed genome assemblies. Given the high heterozygosity of the longan accessions, we performed haplotype-resolved assembly on the nine HiFi-sequenced accessions and ‘Shixia’ combined with chromosome conformation capture (Hi-C) data using hifiasm, while the ONT-sequenced accessions were assembled into diploid genomes using Next-denovo and polished by NextPolish. Overall, the 111 (20 haploid + 91 diploid) genome assemblies had contig N50 ranging from 0.5 to 28.3 Mb (**Fig. 3c**). For the HiFi-sequenced accessions, the initial contigs were anchored to the chromosomes using the Juicer and 3d-DNA pipelines with Hi-C data, and the remaining accessions were anchored to the chromosomes using RagTag^18^ with the T2T assembly as the reference. The sequencing reads from individual accessions were remapped onto the corresponding genomes, with the average mapping ratio at 99.61% (**Table S3**). The completeness of these genomes was estimated to an average of 98.39% by BUSCO. We calculated the assembly quality assessment index (LAI), which had an average of 13.07, meeting the qualification criteria at the reference level or above (**Fig. 3d and Table S3**). The sizes of these longan genome assemblies ranged from 410.3Mb to 493.9 Mb, while each chromosome exhibited similar size through these assemblies (**Fig. 3e, Fig. S5** and **Table S3**). These results demonstrate the high accuracy, completeness and contiguity of the 111 longan genome assemblies. Additionally, the number of gene models were annotated for these genomes ranged from 35,898 to 38,059, with an average of 37,063 protein-coding genes in each assembly (**Table S3**). The gene annotation completeness was further supported by BUSCO (average score of 91.79%; **Fig. S5** and **Table S3**). TEs constituted an average of 56.15% of each genome, ranging from 54.57% to 57.25% across the assembled genomes (**Tables S4**).

**Figure 3.**
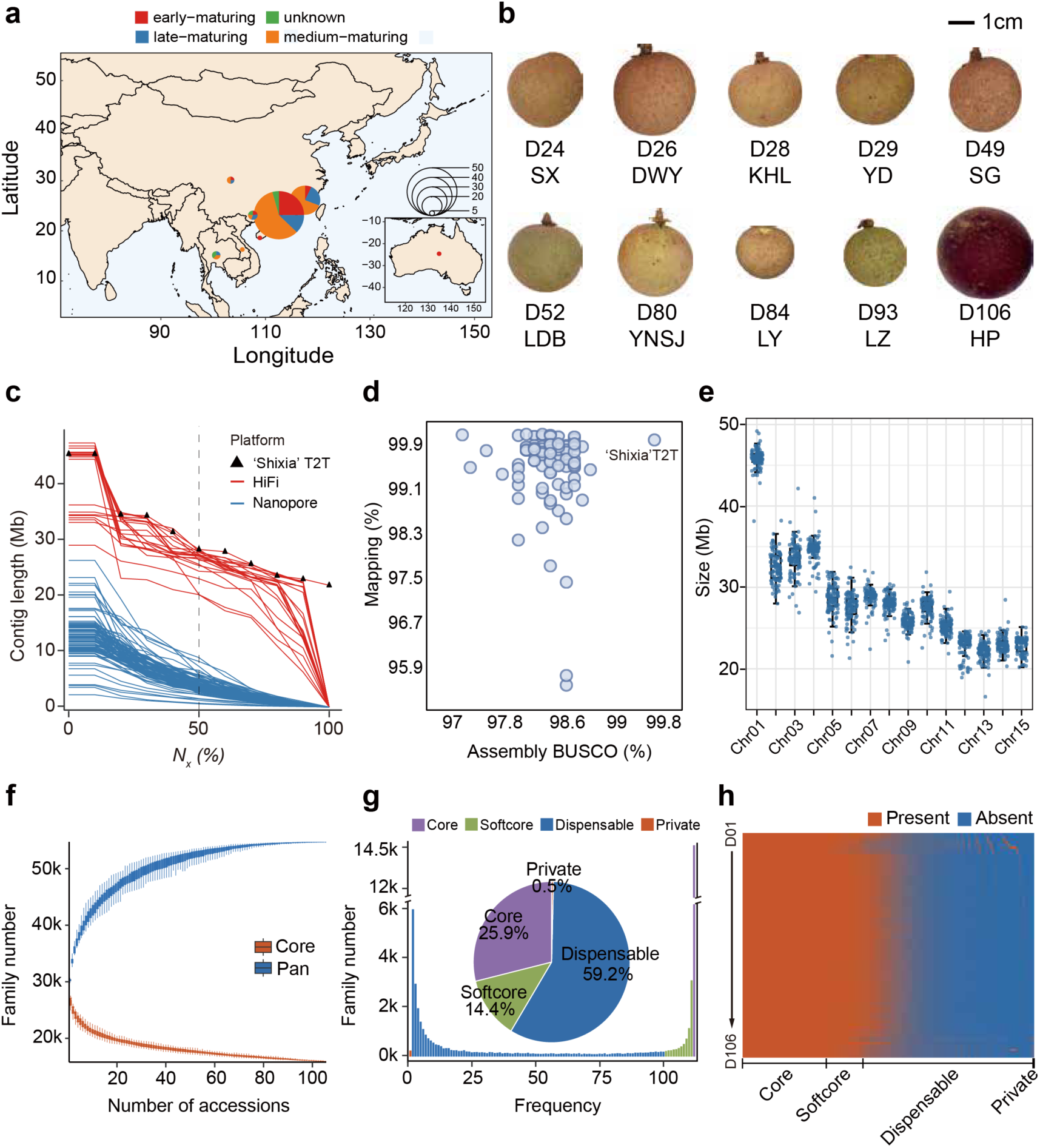
Longan pangenome analysis from 111 chromosome-level genome assemblies. **a**, Geographic distribution of 101 longan accessions sequenced in this study. The color proportion of the circle is proportional to the number of accessions of different types. **b**, Fruit phenotypes of 10 diverse longan accessions for high-quality assemblies. **c**, Comparison of contig Nx size for longan genomic assemblies. The red dashed line marks the contig N50 index. The ’Shixia’ T2T is highlighted by triangle. Accessions sequenced by PacBio HiFi and Oxford Nanopore are colored in red and blue, respectively. **d**, Mapping radio and BUSCO of each longan assembly. **e**, Chromosome sizes of each longan assembly. **f**, The sizes of the pangenome and core-genome models across the 112 longan genomes. **g**, The number and proportion of core (purple), softcore (green), dispensable (blue) and private (red) gene families in the pan-genome. **h**, The PAV (presence/absence variants) of gene families in longan pan-genome.

We next investigated the longan genomic diversity by constructing a pangenome that integrated 111 newly-assembled genome assemblies (20 haploid and 91 diploid) and the annotated ’Shixia’ T2T genome. The pangenome analysis classified all orthologs were into 54,730 gene families (orthogroups). The curve for cumulative number of pan-gene families plateaued as additional longan genomes were iteratively incorporated (**Fig. 3f**), suggesting a nearly-closed pangenome which reflected the broad genetic diversity in our collection of longan accessions. Among the 55,252 gene families, 14,310 (25.9%) were universally present in all 112 genomes (core families), 7,956 (14.4%) were present in 100-111 genomes (softcore families); 32,709 (59.2%) were present in 2-100 genomes, defined as dispensable families, and 277 (0.5%) were present in a single genome, termed private families (**Fig. 3g-h**). The mean nonsynonymous/synonymous substitution (Ka/Ks) ratios for core and softcore genes were significantly lower than those for dispensable and private genes (**Fig. S6a**), suggesting a stronger diversifying selection for the latter two less conserved families. The higher Pfam annotation rates in core and softcore genes suggested that they were more evolutionarily conserved (**Fig. S6b**). Gene ontology (GO) enrichment confirmed that the involvement of core and softcore genes in housekeeping biological processes such as flowering, photosynthesis, hexose mediated signaling (**Fig. S6c**). By contrast, dispensable genes were enriched in cellular response to desiccation, hydrogen peroxide transmembrane transport, and ion channel activity, while private genes were involved in cytokinin dehydrogenase activity (**Fig. S6d**). Together, the pangenome analysis revealed the genetic diversity of longan germplasms as a result of domestication and breeding, and highlighted the functional division among different pan-gene families reflecting their potential contributions to longan evolution.

### A comprehensive catalog of structural variations in longan

Structural variants (SVs) have direct impact on crop traits by disrupting gene functions, thus driving the domestication and breeding. Using ’Shixia’ T2T genome as reference, pairwise whole-genome alignments from 111 chromosome-level assemblies using SyRI^19^ revealed extensive genomic variations and rearrangements among the 112 longan genomes (**Fig. 4a**). A total of high-quality 1,093,145 SVs were identified, including 413,810 deletions, 638,412 insertions, 28,027 translocations, and 12,896 inversions (≥50 bp) (**Fig. 4b, Fig. S7a, Table S6**). The number of identified SVs per genome ranged from 5,034 to 12,362 (**Table S6**), affecting approximately 12.17 Mb to 70.54 Mb of the genomes. The majority of SVs were insertions and deletions located within gene and intergenic regions, affecting a high percentage of Intergenic, upstream and downstream regions, conferring the genomic diversity and regulatory effects on gene functions (**Fig. 4c**). Pan-SV analysis showed that no common SVs were shared by all genomes (**Fig. S7b**) and most SVs were private to individual genomes (**Fig. S7c**), highlighting the substantial genetic divergence among the accessions. Although insertions and deletions were more common, inversions were much longer on average (**Fig. 4b**). The representative insertions and deletions were verified for authenticity using PacBio read mapping, and the large-scale inversions (>1 Mb) were validated using Hi-C heatmaps (**Fig. 4d, Fig. S8a-c**).

**Figure 4.**
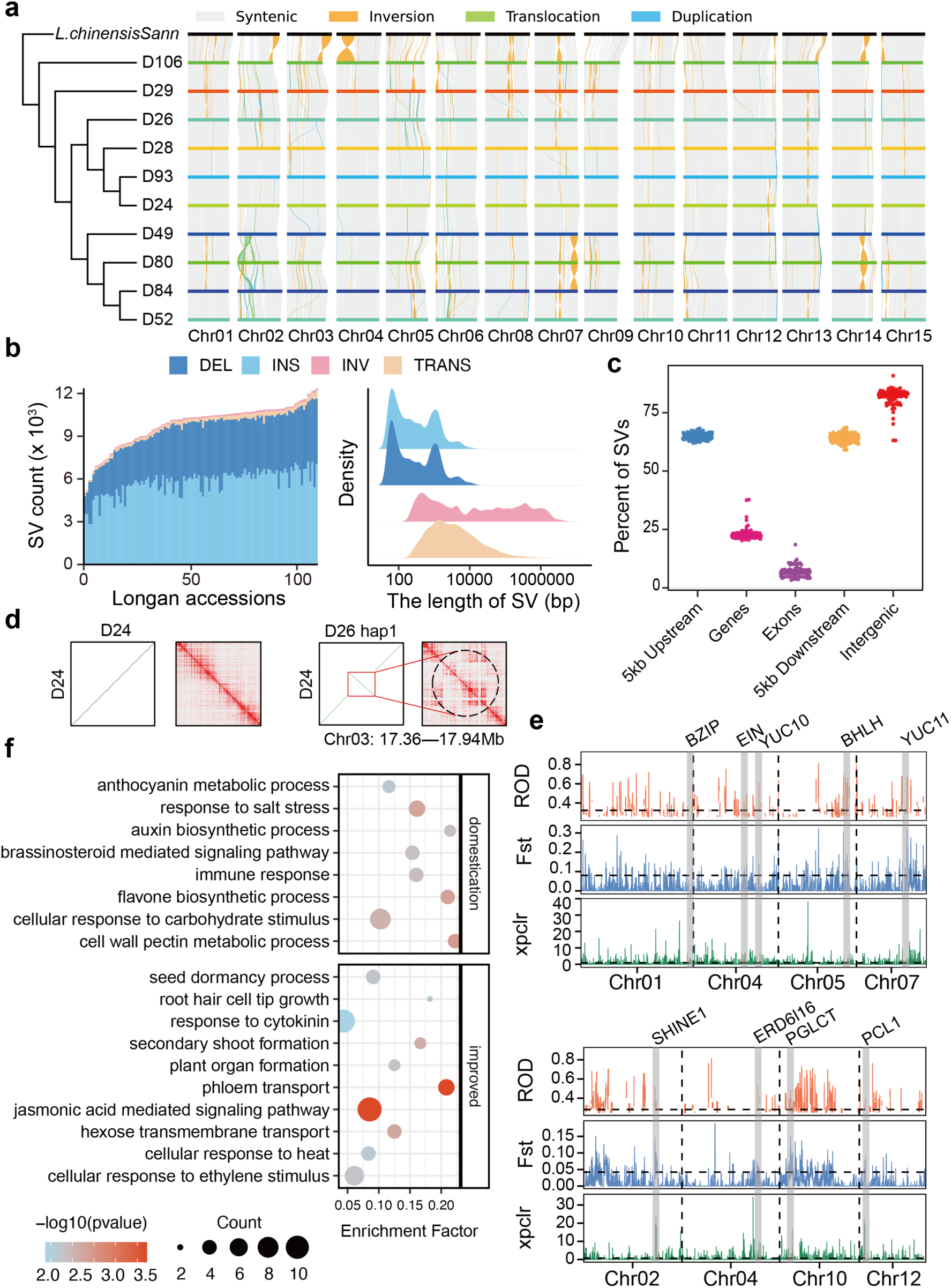
Structural variation analysis reveal signatures of selective sweepings in longan domestication. **a**, SyRI-derived comparative genomic visualization map illustrating the synteny and rearrangements among 10 accessions. **b**, The number and length of different kinds of SVs in longan pan-genome. **c**, Percentage of overlapping features in the pan-genome analysis. The horizontal axis denotes different genomic regions, including upstream, exon, genes, intergenic and downstream. The vertical axis indicates the percentage of overlap of features in the pan-genome for each region. **d**, Example of an inversion event on Chr03 from 17.36 Mb to 17.94 Mb in D26 hap1 (right) validated by Hi-C data, as opposed to ’Shixia’ T2T reference control (left). Left, dot plot showing genome alignment to ’Shixia’ T2T. Right: Hi-C contact heatmap showing Hi-C reads mapping to the ’Shixia’ T2T genome. **e**, Major genomic regions with evidence of selective sweeps during the evolution of landrace accessions from wild accessions and during the evolution of improved accessions from landrace accessions. *Fst*, ROD and XPCLR were used for selection analysis. **f**, GO enrichment analysis of genes in selected regions.

To understand how SVs affect the function of genes during domestication, we further investigated SVs under selection between wild and landrace accessions. Genome-wide analysis of selective sweeps using the fixation index (*Fst*), the reduction in diversity (ROD) and XP-CLR (the cross population composite likelihood ratio test) showed the unbalanced selection of chromosomes during the evolution of landrace accessions from wild accessions and improved accessions from landraces (**Fig. 4e, Fig. S9a-b**). Chr06 (∼10 Mb) and Chr09 (∼9 Mb) had more selection during the evolution of landrace accessions from wild accessions, whereas Chr02 (∼17 Mb), and Chr10 (∼25 Mb) had more regions under selection during the development of cultivated accessions from landraces (**Fig. S9c**). We conducted GO enrichment analysis of the genes within the selected regions. We identified 785 domestication-related genes that have SVs within 10 kb flanking regions, which were enriched in phytohormones biosynthetic and signaling pathway, anthocyanin metabolic process, cell wall pectin metabolic process and stress response. Particularly, the genes homologous to BZIP^20^, bHLH^21^, YUC10 and YUC11^22^, EIN^23,24^ were identified, probably orchestrating morphological adaptations (**Fig. 4f and Table S7**). During the development from landrace to cultivars, the 592 selected genes were mainly involved in sugar transport, formation and development of plant organs, response to cytokinin and ethylene, and other biological regulations (**Fig. 4f and Table S8**), in accordance with deliberate longan breeding for commercially valuable traits (e.g., higher yield, fruit sweetness, and enhanced growth). For example, loss of PCL1 associated with early flowering^25^ in wheat. SHINE1 regulated fruit weight by affecting lipid metabolism^26^ in tomato. PGLCT played a role in the export of starch degradation products^27^ in *Arabidopsis*. In brief, our detailed SV catalog provides crucial insights into the structural genomic diversity of longans, and identified the genomic selective sweeping involving the structural variations that likely underscore the longan domestication and improvement.

### Haplotype-aware landscape of 3D genome organization and gene regulation

Recently, the effect of allele-specific expression (ASE) on fruit quality has received increasing attention^28^, although accurate detection and interaction studies of genetic variation and ASE remains challenging. High-quality haplotype-resolved genome assemblies enable accurate detection of genetic variants and the comparative analysis of three-dimensional chromatin architecture between haplotypes^29^. The high-resolution longan genomic framework allowed us to study the effects of haplotype-specific SVs on ASEs in the context of long traits. In this study, we identified a total of 9,072 ASE genes (ASEGs) across nine representative longan cultivars leveraging their haplotype-resolved genomes, which consistently showed haplotype preferential expression (**Fig. 5a**). Furthermore, GO enrichment analysis revealed that ASE genes in D49 (with thick peel) were enriched in Cutin, suberine and wax biosynthesis, while those in D24 (with high soluble solids content) were enriched in sugar metabolism and transport pathways, implying that ASEGs are closely associated with fruit quality formation (**Fig.5b**). Analysis of haplotype SVs within 5 kb upstream and downstream of all genes showed that the proportion of ASEGs flanked by inter-haplotypic SVs was significantly higher than the genome-wide average (**Fig. 5c**), indicating associations between ASE and sequence divergence between haplotypes. Although the relative abundance of different SV types varied among cultivars, inversions were the predominant SVs associated with ASEGs (**Fig. 5d**), a trend that was different from the overall pangenome (**Fig. 4b**).

**Figure 5.**
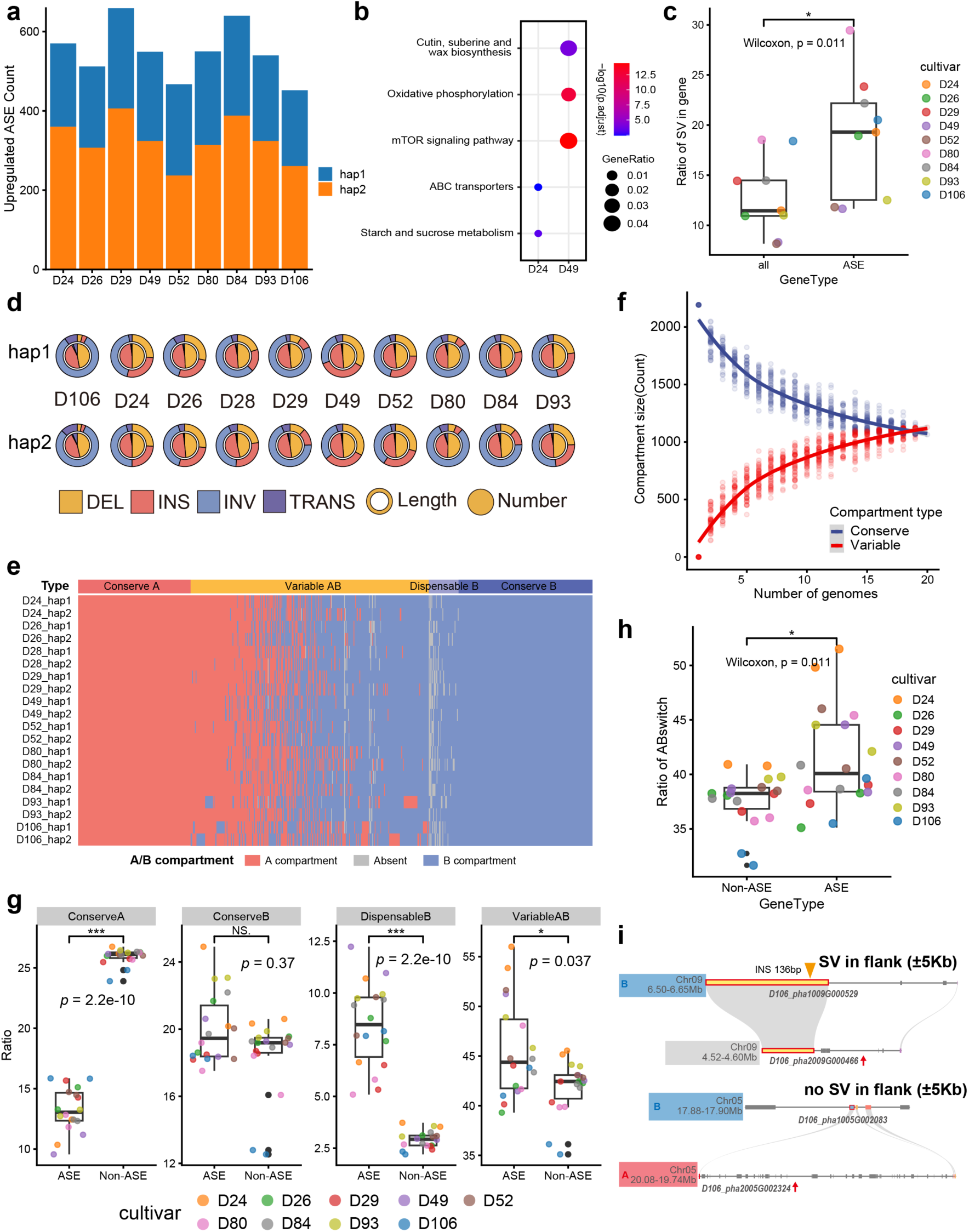
Longan allele-specific expression associated with SV and 3D genome variations. **a**, The upregulated Allele-specific expression (ASE) gene count of 2 haplotypes. **b**, The GO enrichment of ASEGs in D24 and D49. **c**, The ratio of genes (all genes and ASE genes) with SV in flank regions (±5kb). **d**, The distribution of different type of SV in ASEGs in cultivars. **e**, Pan A/B compartment landscape of longans. Absent means one bin with the 0 value of E1. **f**, Pan A/B-genome estimations for compartment types were based on all pairwise whole-genome compartments comparisons across all 20 accessions with sampling max up to 50. **g**, The ratio of ASEGs and Non-ASEGs in different pan-A/B regions. **h**, The ratio of ASEGs and Non-ASEGs in A/B Compartment-Switching Haplotype (CSH) regions. **i**, The example of ASEGs in Dispensable B regions and CSH regions. In **c**,**g**,**h**, n=20, the statistics analysis was based on Wilcoxon signed-rank test, box plots denote the 25th percentile, the median and the 75th percentile, with minimum to maximum whiskers.

Given that 3D chromatin architecture is vital for transcriptional regulation, we explored whether 3D genome organization accounts for ASEs between haplotype genomes by performing a comprehensive pan-3D genome analysis on haplotype-resolved longan assemblies. We observed that 47.9% of A/B chromatin compartments were conserved across the cultivars (47.9% Conserved A/B compartments; **Fig. 5e-f**). This proportion was substantially lower than the 78.5% reported for the soybean pan-3D-genome^30^, likely attributable to the greater genetic diversity among longan cultivars. Notably, 46.1% of the A/B compartments were variable indicative of extensive compartment switching. We found that ASEGs were significantly enriched in Dispensable B and Variable A/B compartments but significantly depleted in Conserved A compartments relative to random distribution (**Fig. 5g**). We wonder whether this enrichment was associated with SVs causing 3D genome variations, and found that Dispensable B compartments exhibited a significantly higher SV density than other compartment types (**Fig. S10a**), suggesting a role for these hypervariable regions in shaping ASEs. Furthermore, the average number of SVs within the 5-kb flanking regions of genes appeared stable, implying that these SVs may regulate ASEs through distal or indirect regulation (**Fig. S10b**). Conversely, the depletion of ASEGs in conserved A compartments was likely attributable to the robustly high transcription activities characteristic of these euchromatic regions, which could mask underlying allelic imbalances (**Fig. S10c**).

We hypothesized that ASEs within the variable A/B compartments could arise from haplotype-variable chromatin conformations. Therefore, we further identified variable A/B regions that undergo compartment switching (Compartment-Switching Haplotype, CSH) between two haplotypes of the same cultivar. The proportion of ASEGs located within CSH regions was significantly higher than that of non-ASEGs (**Fig. 5h**), confirming a strong association between compartment switching and ASEs. Notably, for a considerable number of ASE genes involved in important functions such as anthocyanin biosynthesis, their allele-specific expression occurred in the absence of proximal SVs and was closely associated with different chromatin conformations (**Fig. 5i**). These findings collectively indicated that CSH regions were involved in the regulation of ASE.

### Identification of loci related to fruit and maturity period traits

Genomic variations particularly SVs have major effects on crop traits and evolution, and a non-biased pangenome is required for accurate SV genotyping and trait associations. We constructed a longan graph pangenome using vg pipeline by integrating 58,978 non-redundant SVs (only deletions and insertions) into the ‘Shixia’ T2T genome. This longan pangenome graph, which comprises 1.63 million nodes and 1.64 million edges with a total sequence length of 521.28 Mb, allowed us to rapidly genotype 34,567 SVs across 104 longan accessions by mapping their NGS reads (**Table S1**). We evaluated genotyping accuracy of selected SVs using PCR amplification on 24 longan accessions (**Fig. S11**). The 104 longan cultivars exhibited significant diversity in fruit quality traits, including pericarp thickness, pulp thickness, fruit transverse/longitudinal diameters, total soluble solids (TSS) and seed weight. Among these traits, TSS and seed weight constitute key breeding targets for cultivated longan improvement. Using the variome and phenome data of the 104 cultivars, we performed SNP-GWAS and SV-GWAS which revealed significant SVs and SNPs linked to TSS (**Fig. 6a**), seed weight (**Fig. 6b**). Three SV prominent signal at Chr03:32665147 (P = 5.08 × 10^-7^), Chr13:11531481, Chr13:17791293 (**Fig. 6c**) were significantly associated with TSS. Another QTL loci which lead SNP Chr07:3941545 (P = 7.80 × 10^-7^) and prominent SV signal at Chr14:2579278 (P = 9.69 × 10^-6^) were significantly associated to seed weight (**Fig. 6b**, **Fig. 6d, Table S9**). Among genes in these regions, *DlPLATZ* encoding a homologue of transcription factor, increased seed size and seed weight in Wheat^31^. The SV resides in 58.8 kb upstream of *DlPLATZ* (**Table S10**).

**Figure 6.**
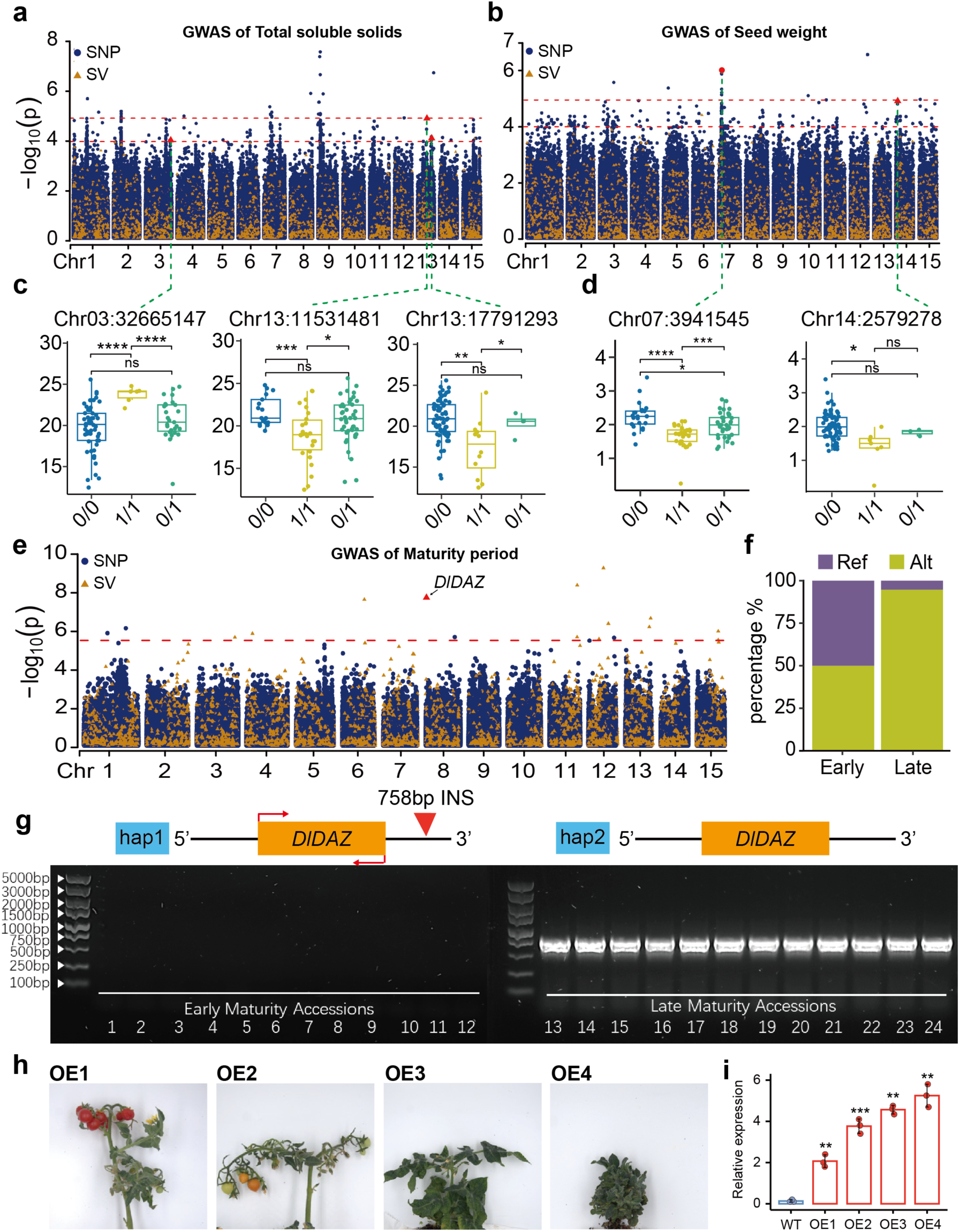
SV-GWAS discovers QTLs and genes associated with longan traits. **a**,**b**, Manhattan plot of SNP- and SV-GWAS performed on total soluble solids and seed weight. **c**,**d** Boxplot of the total soluble solids and seed weight in longan accessions carrying the reference (blue), heterozygote (green) and alternative (yellow) allele. The internal line in each box represents the median and the lower and upper hinges represent the 25th and 75th percentiles, respectively. Statistical significance is examined by a two-sided Student ’s t-test. **e**, Manhattan plot of GWAS performed on maturity period. The red dotted line indicates the significant threshold (p = 2.95 x 10^-6^) from the GEC software. The red arrow indicates the lead variant value (SV) at Chr08:765555 (p = 1.9 x 10^-8^). **f**, Number of longan accessions in different maturity group with the Ref and Alt alleles of the lead SV. **g**, PCR validation of *DlDAZ* (794-bp) two haplotypes in 24 longan accessions. Numbers 1-24 correspond to specific accessions: D04, D15, D22, D32, D38, D40, D51, D55, D60, D81, D82, D83, D11, D16, D17, D35, D36, D62, D63, D65, D85, D86, D90, D91. 1-12 are late maturity varieties, and 13-24 are early maturity varieties. **h**, The over-expression of *DlDAZ* in tomato. **i**, Relative expression levels of *DlDAZ*.

The maturity period critically influences key commercial traits of longan, including sugar accumulation, flavor profile development, as well as harvest scheduling, regional adaptability, and cultivation efficiency. We observed substantial variation in maturation timing across 101 longan accessions. These accessions were classified into early-, mid-, and late-maturing groups, with the D28 ’Kohala’ cultivar representing the early-maturing category, as the first to mature among these accessions. Using the maturity period data of 101 cultivars, SV-GWAS revealed 12 significant SVs (**Table S11**) linked to maturity period, with the most prominent signal being a 758-bp insertion at Chr08:765555 (P = 5.08 × 10^-7^). This SV resided at 539 bp downstream of *DlDAZ*, a gene encoding a C2H2 zinc-finger protein orthologous to *Arabidopsis AtDAZ3*^32^ which functions in controlling sperm cell differentiation^33^.

We further conducted functional validation of this transcription factor. First of all, transient expression in *Nicotiana benthamiana* leaves confirmed the localization of *DIDAZ* in nucleus (**Fig. S12**). Secondly, *DlDAZ* expression levels and the duration of the maturation period were positively correlated (**Fig. 6g**). Finally, stable overexpression of *DlDAZ* in transgenic tomato plants significantly delayed fruit ripening, establishing its conserved role in maturation regulation (**Fig. 6h**). Furthermore, elevated *DlDAZ* expression resulted in a delayed maturation phenotype, and even inhibited the transition from vegetative to reproductive growth by delaying flowering in tomato plants (**Fig. 6h-6i**). The 758-bp insertion within the 3’ UTR at position -539 bp downstream of the *DlDAZ* coding sequence was found in early-maturing accessions, but not in majority of the late-maturing accessions (**Fig. 6f**). Overall, the pangenome graph and GWAS results pave the way for genetic basis of longan traits and lays a theoretical foundation for formulating targeted breeding strategies to improve the commercial value of longan.

## Discussion

The completion of a T2T genome for longan cultivar ’Shixia’, coupled with the construction of a graph-based pangenome encompassing 101 diverse accessions, provides an unprecedented resource for genetic study of this economically important fruit tree. Our study offers a multidimensional view of longan genomic diversity, integrating SVs and 3D chromatin architecture. This comprehensive framework has allowed us to dissect the unique architecture of longan centromeres, uncover the impact of SVs on domestication and breeding, and elucidate a novel mechanism that distal SVs regulate ASE via 3D genome dynamics. Furthermore, it empowers the identification of key genomic loci governing critical agronomic traits, such as fruit maturity period.

The T2T assembly of Shixia has allowed a comprehensive characterization of longan centromeres, revealing a distinctive evolutionary trajectory. In contrast to the centromeres dominated by satellite repeats in *Arabidopsis*^34^, rice^35^, humans^36^, *Vitis*^37^, *N. benthamiana*^38^ and *Brassica*^39^, longan centromeres are primarily composed of intact *Gypsy-CRM* retrotransposons. This architecture aligns with findings in other plant lineages, such as maize^40^, pepper^41^ and lettuce^42^, where *CRM* elements are the most abundant repeats co-localizing with CENH3 nucleosomes. Similarly, in allotetraploid cottons, centromeres are predominantly occupied by *CRM* and *Tekay* elements, as *CRM* plays a critical role in reshaping centromeric structure after polyploidization^43^. Moreover, in Salicaceae, *CRM* elements coupled with LINEs, have extensively colonized *Populus* centromeres^44^. The significant enrichment of intact, CENH3-associating *CRM* elements within the centromeric core, coupled with their pronounced structural preservation compared to their non-centromeric counterparts, strongly suggests that these retrotransposons serve as the primary architectural and functional components of longan centromeres, rather than satellite repeats. The evolutionary dynamics of these centrophilic retrotransposons, however, appear complex and species-specific. While the centromeric retrotransposons are typically younger than those in non-centromeric regions^39,45^, an opposite trend has been observed in *Forsythia suspensa*^46^, highlighting diverse evolutionary histories and turnover rates of centrophilic retrotransposons across plant taxa. We propose that this retrotransposon-dominated centromere model in longan may confer greater epigenetic plasticity, potentially facilitating rapid adaptation to environmental stresses or genomic shocks over the long lifespan of a perennial tree. Validating this hypothesis will require further examination of centromere structures within Sapindaceae family and other perennial trees. These results provide new perspectives for understanding the diversity and evolution of plant centromeres.

Moreover, analysis based on one reference genome may fail to capture pan-species genetic diversity, including structural variants and rare alleles^47^. The comprehensive assessment and utilization of large-scale, representative germplasm resources are critical for longan improvement through identification and capture of genetic variants influencing distinct phenotypes. In this study, high-quality genome assemblies and pan-genome construction of 101 longan accessions enabled the successful identification of several selection signals associated with domestication and improvement traits. The distribution of these loci and genes across populations suggested that selective improvement of agronomic traits was a pivotal driver in longan evolution. Construction of the graph-based genome allowed us to genotype SVs in a large population using short-read resequencing and to perform GWAS. Fruit quality and yield are complex agronomic traits controlled by pleiotropic loci and interconnected genetic networks^48^. We combined SNP and SV genetic markers to conduct association analyses for ten longan fruit traits. Not only did we identify a large number of significant QTL loci and candidate regions, but we also discovered shared genetic signals across different phenotypes (**Fig. S13**), improving the accuracy of genetic analysis and providing a framework for understanding the mechanisms of genetic associations between traits for future longan fruit trait improvement. Indeed, we also find maturity period have higher precision with SV-only markers compared to SNP-only markers (**Fig. 6e**). Among 12 SV-QTLs we pinpointed a 758-bp insertion at downstream of the *DlDAZ* gene, which significantly promotes fruit ripening. This SV site can be utilized as a molecular marker for screening early- and late-maturing longan cultivars in the future. Collectively, our findings facilitate the discovery of domestication genes for longan fruit trait enhancement.

Eukaryotic genes often exhibit allelic imbalance due to a combination of genetic differences between the two chromosome copies^28^. Recently, numerous reports have highlighted the important role of ASE in fruit phenotype^29,49–51^. However, these studies have focused on the ‘nearest gene model’ based on locational proximity, simply assigned the nearest gene as the regulatory target of a genomic variant, neglecting the regulation of ASE by distal variants^52^. The 3D genomics approaches based on the frequency of physical chromatin interactions can predict target genes. We constructed a pan-3D genome using Hi-C data to investigate the relationship between different 3D structures and SVs and ASE genes. We showed that although ASE genes were clearly associated with the dispensable B compartment and the variable A/B compartment, the number of SVs flanking ASE genes did not change significantly. This observation leads us to hypothesize that distal genetic variants may play an important role in the formation of ASE genes, which has been recently supported by growing evidences from other systems. In mammalian cells, extensive folding variability between homologous chromosomes is common, and genetic variation can influence allele-specific expression through 3D structures^53^. For example, haplotype-resolved 3D chromatin architecture in a hybrid pig model revealed that the reorganization of spatial genome between alleles directly causes gene expression imbalance^15^. For example, haplotype-resolved 3D chromatin architecture in a hybrid pig model revealed that the reorganization of spatial genome between alleles directly causes gene expression imbalance^54^. Functional variants can also regulate distant genes by affecting distal regulatory elements (enhancer)^55^. For instance, transvection is a well-established phenomenon in *Drosophila*, where regulatory elements (e.g., enhancers or silencers) on one chromosome allele can act in trans to influence the expression of the homologous allele, as the two alleles are spatially closed^56,57^. However, due to data limitations, we were unable to study the relationship between regulatory elements and ASE genes.

In summary, this study establishes the most comprehensive pan-genome framework for longan, integrating a T2T reference genome with large-scale high-quality genome assemblies. Together, these resources will assist dissecting the biology of longan and the molecular mechanisms by which structural variations regulate complex agronomic traits. The identification and functional validation of key genetic loci associated with fruit quality and maturity timing offer robust molecular targets for precision breeding in longan, highlighting the substantial breeding potential of diverse germplasm resources for improving modern cultivars. As genetic transformation and genome editing technology mature in woody perennial species, the insights generated in this study are poised to accelerate the breeding cycle and facilitate the development of superior longan varieties.

## Methods and Materials

### Plant growth and sampling

A 30-year-old longan tree cultivar named ’Shixia’ from the Fruit Tree Germplasm Repository of Guangdong Province (located at the Institute of Fruit Tree Research at Guangdong Academy of Agricultural Sciences in China) was used for genome sequencing and T2T assembly in this study. From the same repository, 100 additional longan cultivars (**Table 10**) were collected to generate sequencing data for chromosome-scale genome assemblies and pangenome analysis.

### Extraction of DNA and RNA

The extraction of high molecular weight (HMW) genomic DNA was performed using CTAB method. Briefly, 10 g of clean and fresh lettuce leaves were ground with liquid nitrogen and subjected to DNA extraction. The quality of DNA was checked using Qubit 4 (Thermo Fisher Scientific, USA) and Pulsed Field Gel Electrophoresis Systems (Bio-Rad, USA) following manufacturer’s instruction. The isolation of total RNA was performed using TRIzol™ Reagent (Thermo Fisher Scientific, USA) following manufacturer’s instruction. RNA was assessed using RNA Nano 6000 Assay Kit (Agilent Technologies, USA) following manufacturer’s instruction. RNA samples with RNA integrity number (RIN) value > 6.0 were applied for downstream library construction for RNA sequencing.

### DNA library preparation and genome sequencing

For Illumina paired-end sequencing, 5µg of HMW DNA sample was fragmented by sonication to a size of 350 bp and a library was prepared using NEB Next® Ultra™ DNA Library Prep Kit for Illumina (New England Biolabs, USA) following manufacturer’s instruction. The library was sequenced using Illumina Novoseq 6000 platform by Novogene Biotechnologies Inc. (Tianjin, China). For PacBio HiFi sequencing, 15µg of HMW DNA was sheared by g-TUBE (Covaris, USA) and applied for PacBio SMRTbell library using SMRTbell® express template prep kit 2.0 (PacBio, USA). The library was size-selected using BluePippin (Sage Science, USA) with 15 kb as a cutoff. The library was sequenced using PacBio Sequel II system by Novogene Biotechnologies Inc. (Tianjin, China). For Oxford Nanopore sequencing, 1.5µg of HMW DNA was size-selected with 3 kb as a cutoff, and applied for Ligation Sequencing Kit (SQK-LSK109) (Oxford Nanopore Technologies, UK). The DNA ends were FFPE (Formalin-Fixed and Parrffin Embedded) repaired and end-prepped/dA-tailed using NEBNext® Ultra™ II End Repair/dA-Tailing Module (New England Biolabs, USA). Sequencing adapters were ligated onto the prepared ends using the NEBNext® Quick Ligation Module (New England Biolabs, USA). The final DNA library was sequenced using the PromethION sequencer (Oxford Nanopore Technologies, Oxford, UK) by Single-Molecule Sequencing Platform at Novogene Biotechnologies Inc. (Beijing, China).

### Hi-C library preparation and sequencing

The preparation of Hi-C library was performed following the standard protocol described previously with some modifications^16^. Fresh leaves were ground with liquid nitrogen to powder, cross-linked using a 4% formaldehyde solution and ended using 2.5M glycine. The nuclei were suspended in NEB buffer and solubilized using SDS. After digestion with *Dpn* II, the DNA ends of the resultant were ligated with biotin-14-dCTP. The ligated DNA was sheared into 200-600 bp fragments with sonication followed by blunt-end repair and purification through streptavidin C1 magnetic beads pull down. The library was sequenced using Illumina NovaSeq 6000 to obtain 150-bp paired-end reads at Novogene Biotechnologies Inc. (Beijing, China).

### Transcriptome library preparation and sequencing

A total of 3μg of RNA was applied for VAHTS Universal V6 RNA-seq Library Prep Kit for Illumina (Vazyme Biotech, China) following the manufacturer’s instruction. The RNA concentration of the library was measured with Invitrogen™ Qubit™ RNA High Sensitivity (HS), Broad Range (BR), and Extended Range (XR) Assay Kits (Thermo Fisher Scientific, USA), and the concentration was subsequently diluted to 1 ng/µL. After the evaluation of insert size using 2100 Bioanalyzer System (Agilent Technologies, USA), CFX96 Touch Real-Time PCR Detection System (Bio-Rad, USA) was deployed to measure the effective concentration of library (Library effective concentration > 10 nM). The library was sequenced using HiSeq X Ten (Illumina, USA) with paired-end sequencing program (PE150) by Novogene Biotechnologies Inc. (Beijing, China).

### Genome assembly

We selected 101 longan resources of different origins and varieties. Ten accessions were chosen for HiFi sequencing and Hi-C sequencing, and the remaining accessions were only sequenced using ONT. The genome size and heterozygosity were estimated using Jellyfish (v2.2.10)^58^ (k-mer size = 21) and GenomeScope (v2.0)^59^ (max k-mer coverage = 1,000,000) with NGS reads. The preliminary assembly of the HiFi sequenced accessions was completed using hifiasm (0.19.7-r598) with the default parameters. The assembly of the ONT sequenced accessions was performed using NextDenovo^60^ (v2.5.2) and polished with NextPolish^61^ (v1.4.1).

To remove plastid sequences and common contaminants from the genome assemblies, contigs were aligned to mitochondrial and chloroplast genome sequences using minimap2^62^ with the ’-x asm5’ parameter. Contigs with at least 50% base coverage identical to plastid sequences were removed. Additional contaminants were eliminated by aligning contigs to RefSeq bacterial genomes from NCBI using blast-2.13.0+ with the -task megablast parameter. Hi-C sequencing data were then used to anchor contigs with the Juicer (v1.6), 3D-DNA (v180922), and Juicebox (v1.11.08) pipelines for HiFi-sequenced accessions. Assembly validation involved manual checking and orientation tuning of contigs, with any misassemblies corrected in Juicebox. For other ONT-sequenced samples, chromosomes were anchored using RagTag (v2.1.0).

### Genome validation and quality evaluation

For mapping statistics, the NGS short reads were mapped using BWA^63^ (v0.7.17), and the HiFi and ONT long reads were mapped using Minimap2. Then SAMtools^64^ (v1.10) was used to determine the mapping rates and coverage depth. To assess genome completeness, we applied BUSCO^65^ (v5.4.3) for ortholog detection using the embryophyta_odb10. The QV was estimated using Merqury^66^ (v1.3) from HiFi reads. LTR Assembly Index (LAI)^67^ was calculated to evaluate the assembly continuity of LTR retrotransposons.

### Repetitive element and gene annotation

We scanned the genome using RepeatMasker^68^ (v4.1.5) with Repbase^69^ and a de novo repeat database constructed with RepeatModeler^70^ (v2.0.4). RepeatMasker was employed to mask the genome and annotate transposable elements (TEs). Structures of protein-coding genes in the genomes were predicted from three evidences: *ab initio* gene prediction, homology-based gene prediction, and transcripts-based gene prediction. Before gene prediction, the assembled genomes were soft masked using RepeatMasker. MAKER^71^ (v2013-02-16) and BRAKER^72^ (v2.1.6) pipelines were used to perform *ab initio* gene prediction. For homology-based prediction, protein sequences of *Arabidopsis thaliana*, *Carica papaya*, *Dimocarpus longan*, *Nephelium lappaceum*, *Sapindus mukorossi*, *Litchi chinensis* were refined by cd-hit and aligned to our genome assemblies by miniprot^73^ (v0.13-r248). Transcriptome evidence was generated using HISAT2^74^ (v2.2.1), Trinity^75^ (v2.8.4) and StringTie^76^ (v2.2.1) with RNA reads. EVidenceModeler^77^ (v2.1.0) was used to integrate all prediction results to predict gene models. Completeness of the predicted annotation sets were assessed using BUSCO protein mode with the embryophyta_odb10 database. For consistency, the proteome sets were additionally vetted with Omark^78^ (v2.0.3) using default settings with the LUCA.h5 database.

### Hi-C data processing

The HiC-Pro^79^ suite v.3.1.0 was used for valid mapping of the Hi-C reads. Clean Hi-C data from individual accessions were mapped to assembled genome (termed Self mapping in this study) and a common reference genome using bowtie2^80^. During mapping, the reads were first aligned using an end-to-end aligner, and the reads spanning ligation junctions were trimmed at their 3′ end and realigned to the genome. The aligned reads of both fragment mates were then paired in a single paired-end BAM file. Dangling-end reads, same-fragment reads, self-circled reads, self-ligation reads, and other invalid Hi-C reads were discarded from subsequent analyses.

After removing duplications, valid pairs were used to generate raw Hi-C matrix at 10-kb, 20-kb, 50-kb, 100-kb, 200-kb, 500-kb, and 1-Mb resolutions. These matrices were normalized by the iterative correction and eigenvector decomposition (ICE) method. To investigate the contact domains, the valid pairs were converted into .hic format files with the juicer tool pre command; to calculate compartment signals and identify TAD-like or loops, the valid pairs were converted into .cool format files with hicConvertFormat command in HiCExplorer^81^ suite v3.7.2; and to visualize the Hi-C contact map, the valid pairs were converted into .h5 format.

### CENH3 ChIP-seq

Antibody against the full-length CENH3 sequence was prepared and purified using rabbit serum by AtaGenix (Wuhan, China). Then, the ChIP cloning experiment was conducted on chromatin extract from young leaves using the anti-CENH3 antibody as described previously^38^. The ChIP library was amplified with the VAHTS® Universal DNA Library Prep Kit for Illumina V3 (Vazyme ND607) and sequenced on the Illumina NovaSeq 6000 platform to produce 150-bp paired-end reads at Shandong Xiuyue Biotechnology Co., Ltd (Jinan, China).

### Centromere identification and analysis

The centromeres were identified by using CENH3 ChIP-seq reads from each accession according to a previously reported pipeline. Briefly, the paired-end reads were first preprocessed with fastp^82^ (v0.23.2) to remove low-quality bases, and then mapped to each genome assembly using Bowtie2 (v2.5.1) with the following settings: "--very-sensitive --no-mixed --no-discordant -k 10 --maxins 800". All read duplicates and alignments with a mapping quality of less than 10 were discarded. The number of reads overlapped with 25-kb or 250-kb bins of chromosomes were counted. For profiling of CENH3 occupancy, the bamCompare tool from deepTools^83^ (v3.5.1) was used to calculate log2(ChIP/control). Non-B DNA was identified using the Non-B DB v2.0 database^84^.

### Core and dispensable gene family of pangenome construction

The core and dispensable gene sets were classified based on gene family classification from Orthfinder^85^ (v2.5.5). According to clustering results, gene families shared by all samples were defined as core gene families, while gene families missing in one or two genetic resources were defined as soft-core gene families. Gene families missing in more than two resources were categorized as dispensable gene families, and gene families present in only one genetic resource were defined as private gene families. The ratio of non-synonymous to synonymous substitutions (Ka/Ks) was calculated using wgd^86^. Gene function annotation was performed using eggNOG-mapper^87^ (v2.1.12, https://eggnog-mapper.embl.de/). The enrichment test was performed by the ClusterProfiler^88^ package (v3.10.1) in R 4.3.0.

### Structural variation identification

SV identification was performed with Minimap2 and SYRI software. Pairwise genome alignment was executed between the genomes of 111 assemblies. To reduce false positives in SV calling, only SVs such as insertions, deletions, inversions (less than 1 Mb in size), and translocations (greater than 50-bp in length) were retained. Validation of SV with Hi-C data. The paired-end reads from Hi-C are aligned with the corresponding genomic assemblies and the interaction heat maps of the SV-containing regions are manually checked. We also have validated deletion and insertion by aligning PacBio long reads to the ’Shixia’ T2T assembly, followed by checking the read mapping at the breakpoints to identify the split-read evidence, using IGV (Integrative Genomics Viewer) visualization.

### Construction of the graph-based pangenome and pan-SVs genotyping

To integrate linear reference genomes with large-scale genome variants for the construction of comprehensive variant maps, the identified SVs (only insertions and deletions) in 111 de novo assembled genomes against the ’Shixia’ T2T reference were merged using SURVIVOR^89^ (v1.0.7) with the specific parameters "1000 0 1 1 1 0 50". The graphed pangenome was then constructed using vg^90^ pipeline (v1.46.0) with the parameters " autoindex --workflow giraffe".

For genotype SVs, Illumina short reads from each accession were aligned to the graph-based super-pangenome using giraffe tool from vg pipeline with default parameters. Poor alignments with quality scores below five were removed using vg with the parameters "pack -Q 5 -s". SV genotyping was then performed using vg calls with default parameters.

### ASE analysis

Syntenic gene blocks between the two haplotype genomes were identified using MCScanX (v1.0.0)^91^, with gene pairs located within large syntenic blocks classified as alleles. The assemblies and annotations of both haplotype genomes were subsequently merged to construct a metagenome. Clean transcriptomic reads were then mapped to the metagenome using STAR (v2.7.11b)^92^, with the following parameters: -- outMultimapperOrder Random, --outSAMtype BAM Unsorted, --quantMode GeneCounts, --alignIntronMax 6000, --outSJfilterReads Unique, and --outFilterMismatchNmax 1. The resulting BAM files were sorted and PCR duplicates were removed using SAMtools and Picard. The number of reads mapped to genes were counted using FeatureCounts^93^. Genes exhibiting expression levels >1 TPM in at least one haplotype genome were considered allele-specific expressed (ASE) genes^94^.

### Pan-3D genome architecture identification

The E1 values from the eigenvector decomposition on Hi-C contact maps were used to indicate the A/B compartment status. We used cooltools (v0.7.0)^95^ with the parameter “cooltools call-compartments” to obtain the E1, E2, and E3 values based on a 1mb resolution Hi-C contact matrix. Because E2 or E3 values sometimes reflect A/B compartments, we manually checked the E1, E2, and E3 tracks with gene density and the plaid pattern in the Hi-C contact maps along each chromosome and obtained the final “E1” list. The direction of the eigenvalues is set to A or B based on their association with gene expression (CPM). Compartments were divided into Switch A/B and Stable A/B, compared to the upstream bins (Switch A/B means the compartment is different from the upstream contiguous bin). To transfer the target genome A/B compartment to the reference genome, minimap v2.28, transanno v0.4.5 (https://github.com/informationsea/transanno) and Crossmap^96^ v0.7.0 were used in pipeline. Then we calculated the weighted average balanced E1 to identify the A/B compartment. According to each bin compartment status, all the bins on the reference genome were divided into five types: Conserve A or B (A or B compartment in all samples), Dispensable A or B (A or B compartment or absence (E1=0 or NA) in all samples, with different compartment was not allowed) and Variable A/B.

### Genome-wide association study

GWAS was performed on SNPs and SVs for 104 accessions using mixed linear modelling with EMMAX^97^ (v.beta-07Mar2010). The first ten principal components and an IBS kinship matrix derived from all SNPs or SVs computed by EMMAX were included as covariates to control for population structure. The number of effective SVs was determined using the Genetic type 1 Error Calculator (GEC)^98^ (v0.2). The results were presented using the CMplot^99^ package in the R environment.

### Tomato transformation and subcellular localization analysis

A 794-bp fragment of the *DlDAZ* coding region was amplified using the PCR primers (listed in Table S12) and introduced to generate the overexpressed plasmid. *A. tumefaciens* EHA105 containing the recombinant vectors pBWA(V)HS-*DlDAZ*-osGFP was used to transform tomato (microTom)^100^. Transgenic tomato plants were selected on MS medium supplemented with 30mg L^-1^ hygromycin and their identity was confirmed through PCR. Subsequently, the T3 generation of transgenic plants was utilized for phenotypic analysis of the candidate gene *DlDAZ*.

The recombinant plasmids pCAMBIA1300-*DlDAZ*-GFP (35S:*DlDAZ*-GFP) marker vector and empty control vector were transformed into *A. tumefaciens* strain GV3101. Agrobacterium mediated transient expression assays in tobacco (*Nicotiana benthamiana*) using the method reported previously^101^. Fluorescence images of transiently infected tobacco leaf subepidermal cells were captured utilizing laser scanning confocal microscopy (catalogue number LSM710, Carl Zeiss).

### Genotype analysis of early- and late-maturity Longan cultivars

Leaf and fruit samples of Longan cultivars were collected for genomic DNA extraction from leaf samples to analyze the genotype of the SV insertion (s2410094), PCR amplification was carried out using 2 x Rapid Taq Plus Master Mix (catalogue number P223-01, Vazyme). Primers (as detailed in Table S12) were used to generate a 2238-bp product corresponding to the 972-bp insertion Chr12:16708690. For RNA extraction from fruit pulp samples to analyze the genotype of *DlDAZ*, RT-PCR was carried out using HiScript IV All-in-One Ultra RT SuperMix (catalogue number R433-01, Vazyme) to generate a 794-bp gene product. The PCR products were visualized on a 1% agarose gel.

## Statistical analysis

For relative expression and sucrose content comparison, t tests were used for significance testing.

## Author contributions

LG conceived and designed the project. JXW, JS, KW, XFW, SC performed genomic analysis. JW, JBW and JYL performed experiments. JW and JL provided plant materials for sequencing. JXW, LG, JS, KW and XLR wrote and revised the manuscript. All authors read and approved the final version of the manuscript.

## Acknowledgements

This work was supported by the Shandong Provincial Natural Science Foundation (SYS202206), Key R&D Program of Shandong Province (ZR202211070163), Taishan Scholars Program and Natural Science Foundation for Distinguished Young Scholars of Shandong Province (ZR2023JQ010). The authors would like to thank Bioinformatic Platform in Peking University IAAS for providing high-performance computing resources.

## References

1. Wu, Y., Yi, G., Zhou, B., Zeng, J. & Huang, Y. The advancement of research on litchi and longan germplasm resources in China. Scientia Horticulturae 114, 143–150 (2007).

2. Zeng, S., Wang, K., Liu, X., Hu, Z. & Zhao, L. Potential of longan (Dimocarpus longan Lour.) in functional food: A review of molecular mechanism-directing health benefit properties. Food Chemistry 437, 137812 (2024).

3. Giovannoni, J. M OLECULAR B IOLOGY OF F RUIT M ATURATION AND R IPENING. Annu. Rev. Plant. Physiol. Plant. Mol. Biol. 52, 725–749 (2001).

4. Lin, Y. et al. Genome-wide sequencing of longan ( *Dimocarpus longan* Lour.) provides insights into molecular basis of its polyphenol-rich characteristics. GigaScience 6, gix023 (2017).

5. Wang, J. et al. Genomic insights into longan evolution from a chromosome-level genome assembly and population genomics of longan accessions.

6. Liao, L. et al. Unraveling a genetic roadmap for improved taste in the domesticated apple. Molecular Plant 14, 1454–1471 (2021).

7. Weischenfeldt, J., Symmons, O., Spitz, F. & Korbel, J. O. Phenotypic impact of genomic structural variation: insights from and for human disease. Nat Rev Genet 14, 125–138 (2013).

8. Qin, P. et al. Pan-genome analysis of 33 genetically diverse rice accessions reveals hidden genomic variations. Cell 184, 3542–3558.e16 (2021).

9. Li, B. et al. A gap-free reference genome reveals structural variations associated with flowering time in rapeseed ( *Brassica napus* ). Horticulture Research 10, uhad171 (2023).

10. Chen, J. et al. Pangenome analysis reveals genomic variations associated with domestication traits in broomcorn millet. Nat Genet 55, 2243–2254 (2023).

11. Chen, W. et al. Graph pangenome reveals the regulation of malate content in blood-fleshed peach by NAC transcription factors. Genome Biol 26, 7 (2025).

12. Schreiber, M., Jayakodi, M., Stein, N. & Mascher, M. Plant pangenomes for crop improvement, biodiversity and evolution. Nat Rev Genet 25, 563–577 (2024).

13. Cheng, H., Concepcion, G. T., Feng, X., Zhang, H. & Li, H. Haplotype-resolved de novo assembly using phased assembly graphs with hifiasm. Nature Methods 18, 170–175 (2021).

14. Durand, N. C. et al. Juicer Provides a One-Click System for Analyzing Loop-Resolution Hi-C Experiments. Cell Systems 3, 95–98 (2016).

15. Dudchenko, O. et al. De novo assembly of the Aedes aegypti genome using Hi-C yields chromosome-length scaffolds. Science 356, 92–95 (2017).

16. Durand, N. C. et al. Juicebox Provides a Visualization System for Hi-C Contact Maps with Unlimited Zoom. Cell Syst 3, 99–101 (2016).

17. Smeds, L. et al. Non-canonical DNA in human and other ape telomere-to-telomere genomes. Nucleic Acids Res 53, gkaf298 (2025).

18. Alonge, M. et al. Automated assembly scaffolding using RagTag elevates a new tomato system for high-throughput genome editing. Genome Biol 23, 258 (2022).

19. Goel, M., Sun, H., Jiao, W.-B. & Schneeberger, K. SyRI: finding genomic rearrangements and local sequence differences from whole-genome assemblies. Genome Biol 20, 277 (2019).

20. Lu, Z. et al. SpbZIP60 confers cadmium tolerance by strengthening the root cell wall compartmentalization in Sedum plumbizincicola. J Hazard Mater 480, 135936 (2024).

21. Xu, Y. et al. Gibberellin signaling regulates pectin biosynthesis in Arabidopsis. Nat Commun 16, 4065 (2025).

22. Hu, Y. et al. Gibberellins play an essential role in late embryogenesis of Arabidopsis. Nat Plants 4, 289–298 (2018).

23. Yanagisawa, S., Yoo, S.-D. & Sheen, J. Differential regulation of EIN3 stability by glucose and ethylene signalling in plants. Nature 425, 521–525 (2003).

24. Guo, R. et al. Arabidopsis EIN2 represses ABA responses during germination and early seedling growth by inactivating HLS1 protein independently of the canonical ethylene pathway. Plant J 115, 1514–1527 (2023).

25. Mizuno, N. et al. Loss-of-Function Mutations in Three Homoeologous PHYTOCLOCK 1 Genes in Common Wheat Are Associated with the Extra-Early Flowering Phenotype. PLoS One 11, e0165618 (2016).

26. Li, Q. et al. Differential expression of SlKLUH controlling fruit and seed weight is associated with changes in lipid metabolism and photosynthesis-related genes. J Exp Bot 72, 1225–1244 (2021).

27. Cho, M.-H. et al. Role of the plastidic glucose translocator in the export of starch degradation products from the chloroplasts in Arabidopsis thaliana. New Phytol 190, 101–112 (2011).

28. Sun, X. et al. Phased diploid genome assemblies and pan-genomes provide insights into the genetic history of apple domestication. Nat Genet 52, 1423–1432 (2020).

29. Li, Q. et al. Haplotype-resolved T2T genome assemblies and pangenome graph of pear reveal diverse patterns of allele-specific expression and the genomic basis of fruit quality traits. Plant Commun 5, 101000 (2024).

30. Pan-3D genome analysis reveals structural and functional differentiation of soybean genomes - PubMed. https://pubmed.ncbi.nlm.nih.gov/36658660/.

31. Fu, Y. et al. Identification and Characterization of PLATZ Transcription Factors in Wheat. Int J Mol Sci 21, 8934 (2020).

32. Liu, Y., Khan, A. R. & Gan, Y. C2H2 Zinc Finger Proteins Response to Abiotic Stress in Plants. Int J Mol Sci 23, 2730 (2022).

33. Jia, H.-H. et al. Genome-wide identification of the C2H2-Zinc finger gene family and functional validation of CsZFP7 in citrus nucellar embryogenesis. Plant Reprod 36, 287–300 (2023).

34. Naish, M. et al. The genetic and epigenetic landscape of the Arabidopsis centromeres. Science 374, eabi7489 (2021).

35. Cheng, Z. et al. Functional rice centromeres are marked by a satellite repeat and a centromere-specific retrotransposon. Plant Cell 14, 1691–1704 (2002).

36. Altemose, N. et al. Complete genomic and epigenetic maps of human centromeres. Science 376, eabl4178 (2022).

37. Guo, L. et al. Super pangenome of Vitis empowers identification of downy mildew resistance genes for grapevine improvement. Nat Genet 57, 741–753 (2025).

38. Chen, W. et al. The complete genome assembly of Nicotiana benthamiana reveals the genetic and epigenetic landscape of centromeres. Nat Plants 10, 1928–1943 (2024).

39. Chen, W. et al. Pan-centromere landscape and dynamic evolution in Brassica plants. Nat Plants https://doi.org/10.1038/s41477-025-02131-5 (2025) doi:10.1038/s41477-025-02131-5.

40. Chen, J. et al. A complete telomere-to-telomere assembly of the maize genome. Nat Genet 55, 1221–1231 (2023).

41. Chen, W. et al. Two telomere-to-telomere gapless genomes reveal insights into Capsicum evolution and capsaicinoid biosynthesis. Nat Commun 15, 4295 (2024).

42. Wang, K. et al. The complete telomere-to-telomere genome assembly of lettuce. Plant Communications 5, 101011 (2024).

43. Chang, X. et al. High-quality Gossypium hirsutum and Gossypium barbadense genome assemblies reveal the landscape and evolution of centromeres. Plant Commun 5, 100722 (2024).

44. Wang, Y. et al. Four near-complete genome assemblies reveal the landscape and evolution of centromeres in Salicaceae. Genome Biol 26, 111 (2025).

45. Chen, W. et al. Two telomere-to-telomere gapless genomes reveal insights into Capsicum evolution and capsaicinoid biosynthesis. Nat Commun 15, 4295 (2024).

46. Cui, J. et al. The gap-free genome of Forsythia suspensa illuminates the intricate landscape of centromeres. Hortic Res 11, uhae185 (2024).

47. Guo, D. et al. A pangenome reference of wild and cultivated rice. Nature 642, 662–671 (2025).

48. Zhou, Y. Dissecting the genetic basis of agronomic traits by multi-trait GWAS and genetic networks in maize (Zea mays L.). (2025).

49. Sun, M. et al. Haplotype-resolved, gap-free genome assemblies provide insights into the divergence between Asian and European pears. Nat Genet 57, 2040–2051 (2025).

50. Tian, Y. et al. Transposon insertions regulate genome-wide allele-specific expression and underpin flower colour variations in apple (Malus spp.). Plant Biotechnol J 20, 1285–1297 (2022).

51. Liu, C. et al. The OsWRKY72-OsAAT30/OsGSTU26 module mediates reactive oxygen species scavenging to drive heterosis for salt tolerance in hybrid rice. J Integr Plant Biol 66, 709–730 (2024).

52. Shen, L. et al. Integrative 3D genomics with multi-omics analysis and functional validation of genetic regulatory mechanisms of abdominal fat deposition in chickens. Nat Commun 15, 9274 (2024).

53. Irastorza-Azcarate, I. et al. Extensive folding variability between homologous chromosomes in mammalian cells. bioRxiv 2024.05.08.591087 (2024) doi:10.1101/2024.05.08.591087.

54. Lin, Y. et al. Haplotype-resolved 3D chromatin architecture of the hybrid pig. Genome Res 34, 310–325 (2024).

55. Zunjarrao, S. & Gambetta, M. C. Principles of long-range gene regulation. Curr Opin Genet Dev 91, 102323 (2025).

56. Galouzis, C. C. & Prud’homme, B. Transvection regulates the sex-biased expression of a fly X-linked gene. Science 371, 396–400 (2021).

57. Urban, E. A. et al. Activating and repressing gene expression between chromosomes during stochastic fate specification. Cell Rep 42, 111910 (2023).

58. Marçais, G. & Kingsford, C. A fast, lock-free approach for efficient parallel counting of occurrences of k-mers. Bioinformatics 27, 764–770 (2011).

59. Ranallo-Benavidez, T. R., Jaron, K. S. & Schatz, M. C. GenomeScope 2.0 and Smudgeplot for reference-free profiling of polyploid genomes. Nat Commun 11, 1432 (2020).

60. Hu, J. et al. NextDenovo: an efficient error correction and accurate assembly tool for noisy long reads. Genome Biol 25, 107 (2024).

61. Hu, J., Fan, J., Sun, Z. & Liu, S. NextPolish: a fast and efficient genome polishing tool for long-read assembly. Bioinformatics 36, 2253–2255 (2020).

62. Li, H. Minimap2: pairwise alignment for nucleotide sequences. Bioinformatics 34, 3094– 3100 (2018).

63. Li, H. & Durbin, R. Fast and accurate short read alignment with Burrows-Wheeler transform. Bioinformatics 25, 1754–1760 (2009).

64. Li, H. et al. The Sequence Alignment/Map format and SAMtools. Bioinformatics 25, 2078– 2079 (2009).

65. Simão, F. A., Waterhouse, R. M., Ioannidis, P., Kriventseva, E. V. & Zdobnov, E. M. BUSCO: assessing genome assembly and annotation completeness with single-copy orthologs. Bioinformatics 31, 3210–3212 (2015).

66. Rhie, A., Walenz, B. P., Koren, S. & Phillippy, A. M. Merqury: reference-free quality, completeness, and phasing assessment for genome assemblies. Genome Biol 21, 245 (2020).

67. Ou, S., Chen, J. & Jiang, N. Assessing genome assembly quality using the LTR Assembly Index (LAI). Nucleic Acids Res 46, e126 (2018).

68. Tarailo-Graovac, M. & Chen, N. Using RepeatMasker to identify repetitive elements in genomic sequences. Curr Protoc Bioinformatics **Chapter** 4, 4.10.1–4.10.14 (2009).

69. Bao, W., Kojima, K. K. & Kohany, O. Repbase Update, a database of repetitive elements in eukaryotic genomes. Mob DNA 6, 11 (2015).

70. Flynn, J. M. et al. RepeatModeler2 for automated genomic discovery of transposable element families. Proc Natl Acad Sci U S A 117, 9451–9457 (2020).

71. Cantarel, B. L. et al. MAKER: an easy-to-use annotation pipeline designed for emerging model organism genomes. Genome Res 18, 188–196 (2008).

72. Hoff, K. J., Lomsadze, A., Borodovsky, M. & Stanke, M. Whole-Genome Annotation with BRAKER. Methods Mol Biol 1962, 65–95 (2019).

73. Li, H. Protein-to-genome alignment with miniprot. Bioinformatics 39, btad014 (2023).

74. Kim, D., Paggi, J. M., Park, C., Bennett, C. & Salzberg, S. L. Graph-based genome alignment and genotyping with HISAT2 and HISAT-genotype. Nat Biotechnol 37, 907–915 (2019).

75. Grabherr, M. G. et al. Full-length transcriptome assembly from RNA-Seq data without a reference genome. Nat Biotechnol 29, 644–652 (2011).

76. Shumate, A., Wong, B., Pertea, G. & Pertea, M. Improved transcriptome assembly using a hybrid of long and short reads with StringTie. PLoS Comput Biol 18, e1009730 (2022).

77. Haas, B. J. et al. Automated eukaryotic gene structure annotation using EVidenceModeler and the Program to Assemble Spliced Alignments. Genome Biol 9, R7 (2008).

78. Nevers, Y. et al. Quality assessment of gene repertoire annotations with OMArk. Nat Biotechnol 43, 124–133 (2025).

79. Servant, N., et al. HiC-Pro: an optimized and flexible pipeline for Hi-C data processing. Genome Biol 16, 259 (2015).

80. Langmead, B. & Salzberg, S. L. Fast gapped-read alignment with Bowtie 2. Nat Methods 9, 357–359 (2012).

81. Ramírez, F. et al. High-resolution TADs reveal DNA sequences underlying genome organization in flies. Nat Commun 9, 189 (2018).

82. Chen, S., Zhou, Y., Chen, Y. & Gu, J. fastp: an ultra-fast all-in-one FASTQ preprocessor. Bioinformatics 34, i884–i890 (2018).

83. Ramírez, F. et al. deepTools2: a next generation web server for deep-sequencing data analysis. Nucleic Acids Res 44, W160–165 (2016).

84. Cer, R. Z. et al. Non-B DB v2.0: a database of predicted non-B DNA-forming motifs and its associated tools. Nucleic Acids Res 41, D94–D100 (2013).

85. Emms, D. M. & Kelly, S. OrthoFinder: phylogenetic orthology inference for comparative genomics. Genome Biol 20, 238 (2019).

86. Chen, H. & Zwaenepoel, A. Inference of Ancient Polyploidy from Genomic Data. Methods Mol Biol 2545, 3–18 (2023).

87. Cantalapiedra, C. P., Hernández-Plaza, A., Letunic, I., Bork, P. & Huerta-Cepas, J. eggNOG-mapper v2: Functional Annotation, Orthology Assignments, and Domain Prediction at the Metagenomic Scale. Mol Biol Evol 38, 5825–5829 (2021).

88. Yu, G., Wang, L.-G., Han, Y. & He, Q.-Y. clusterProfiler: an R package for comparing biological themes among gene clusters. OMICS 16, 284–287 (2012).

89. Jeffares, D. C. et al. Transient structural variations have strong effects on quantitative traits and reproductive isolation in fission yeast. Nat Commun 8, 14061 (2017).

90. Hickey, G. et al. Genotyping structural variants in pangenome graphs using the vg toolkit. Genome Biol 21, 35 (2020).

91. Wang, Y. et al. MCScanX: a toolkit for detection and evolutionary analysis of gene synteny and collinearity. Nucleic Acids Res 40, e49 (2012).

92. Dobin, A. et al. STAR: ultrafast universal RNA-seq aligner. Bioinformatics 29, 15–21 (2013).

93. Liao, Y., Smyth, G. K. & Shi, W. featureCounts: an efficient general purpose program for assigning sequence reads to genomic features. Bioinformatics 30, 923–930 (2014).

94. Han, X. et al. Two haplotype-resolved, gap-free genome assemblies for Actinidia latifolia and Actinidia chinensis shed light on the regulatory mechanisms of vitamin C and sucrose metabolism in kiwifruit. Mol Plant 16, 452–470 (2023).

95. Open2C, et al. Cooltools: Enabling high-resolution Hi-C analysis in Python. PLoS Comput Biol 20, e1012067 (2024).

96. Zhao, H. et al. CrossMap: a versatile tool for coordinate conversion between genome assemblies. Bioinformatics 30, 1006–1007 (2014).

97. Kang, H. M. et al. Variance component model to account for sample structure in genome-wide association studies. Nat Genet 42, 348–354 (2010).

98. Li, M.-X., Yeung, J. M. Y., Cherny, S. S. & Sham, P. C. Evaluating the effective numbers of independent tests and significant p-value thresholds in commercial genotyping arrays and public imputation reference datasets. Hum Genet 131, 747–756 (2012).

99. Yin, L. et al. rMVP: A Memory-efficient, Visualization-enhanced, and Parallel-accelerated Tool for Genome-wide Association Study. Genomics Proteomics Bioinformatics 19, 619–628 (2021).

100. Juan, J. X. et al. Agrobacterium-mediated transformation of tomato with the ICE1 transcription factor gene. Genet Mol Res 14, 597–608 (2015).

101. Zhao, K. et al. Pangenome analysis reveals structural variation associated with seed size and weight traits in peanut. Nat Genet 57, 1250–1261 (2025).

