## Supplemental Figures for "Genomic and pan-genomic insight into longan domestication and improvement"

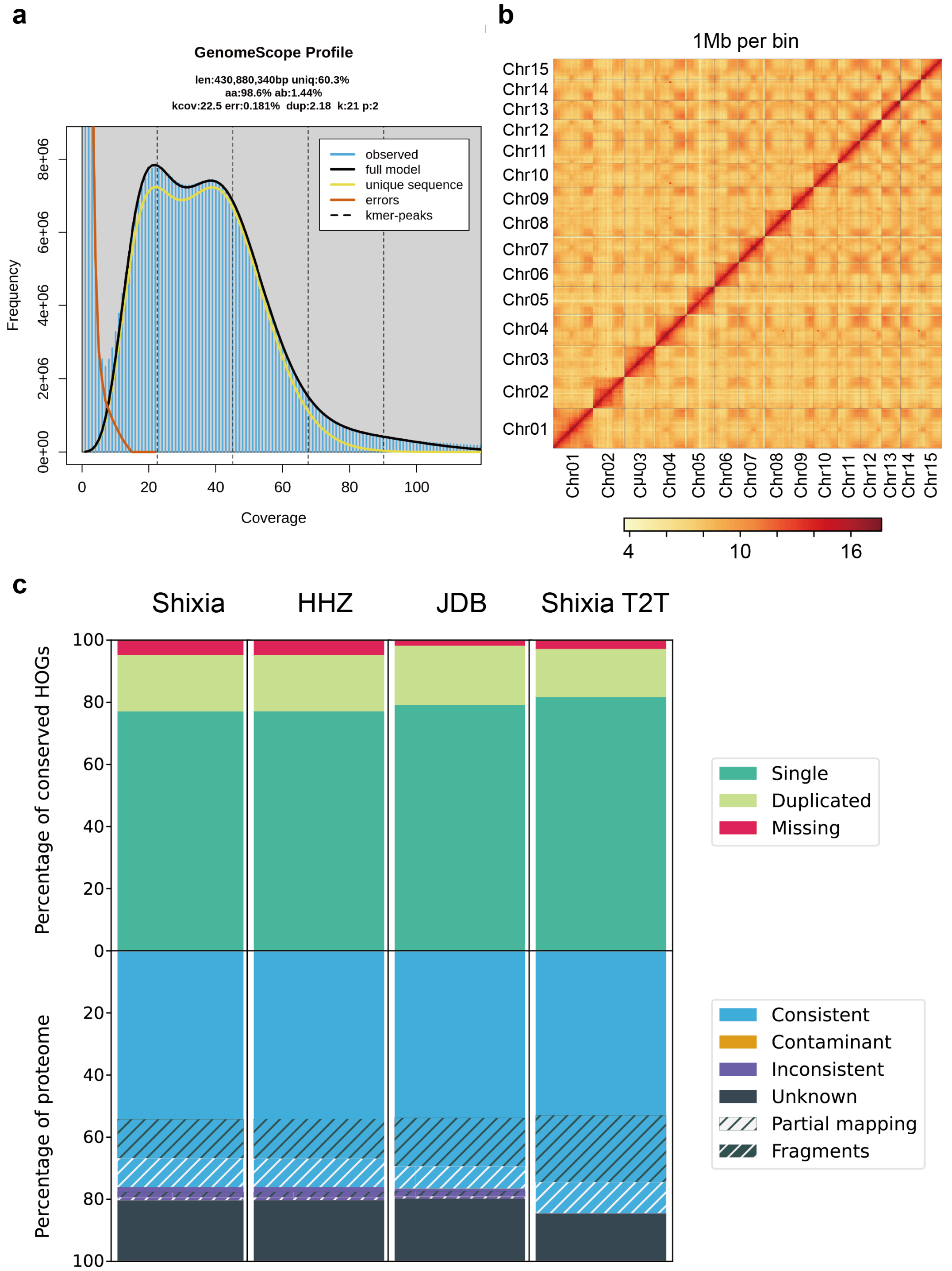


**Supplemental Figure 1. | Genome assembly and annotation of 'Shixia' T2T genome. a**, K-mer frequency analysis was performed using Illumina short reads (55 x coverage) with Jellyfish and GenomeScope2. **b**, Hi-C interaction heatmaps of 'Shixia' T2T genome. The color bars are scaled according to the interaction intensity for each 1 Mb bin. **c**, Comparison of the annotated proteomes from 'Shixia' from NCBI (ASM2045787v1), 'Hong hezi', 'JDB' and 'Shixia' T2T genome.

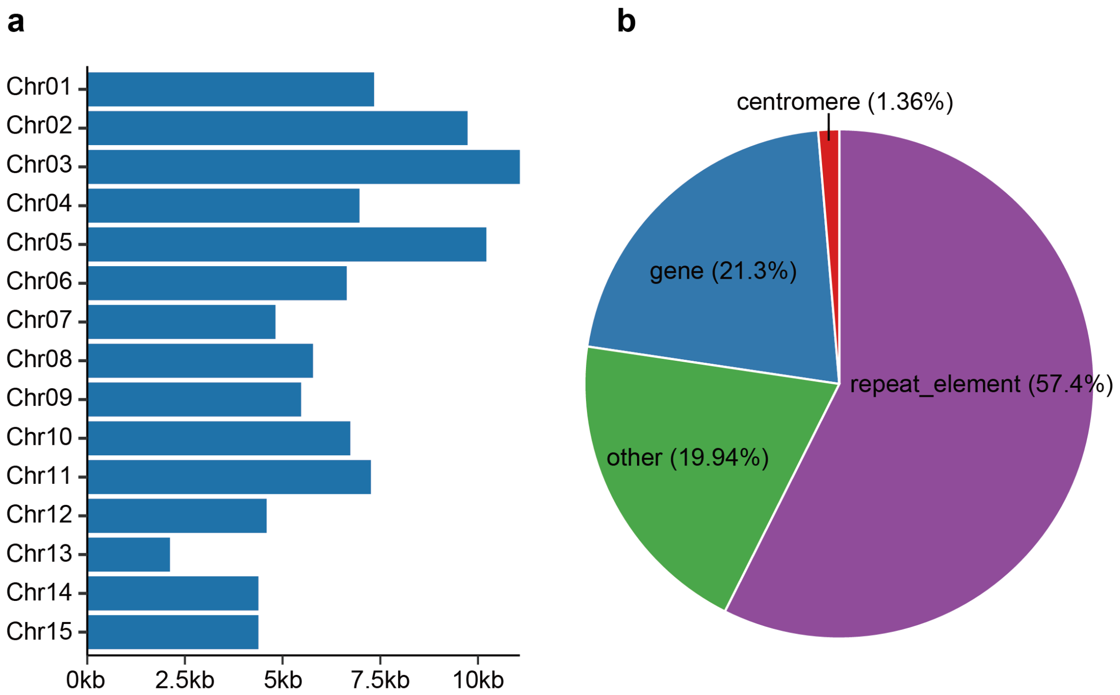


**Supplemental Figure 2.** | **Characterization of newly assembled sequence regions in 'Shixia' T2T genome. a**, Statistics on the length of the newly assembled sequence regions by each chromosome. **b**, Statistics on the composition of the newly assembled sequence regions.

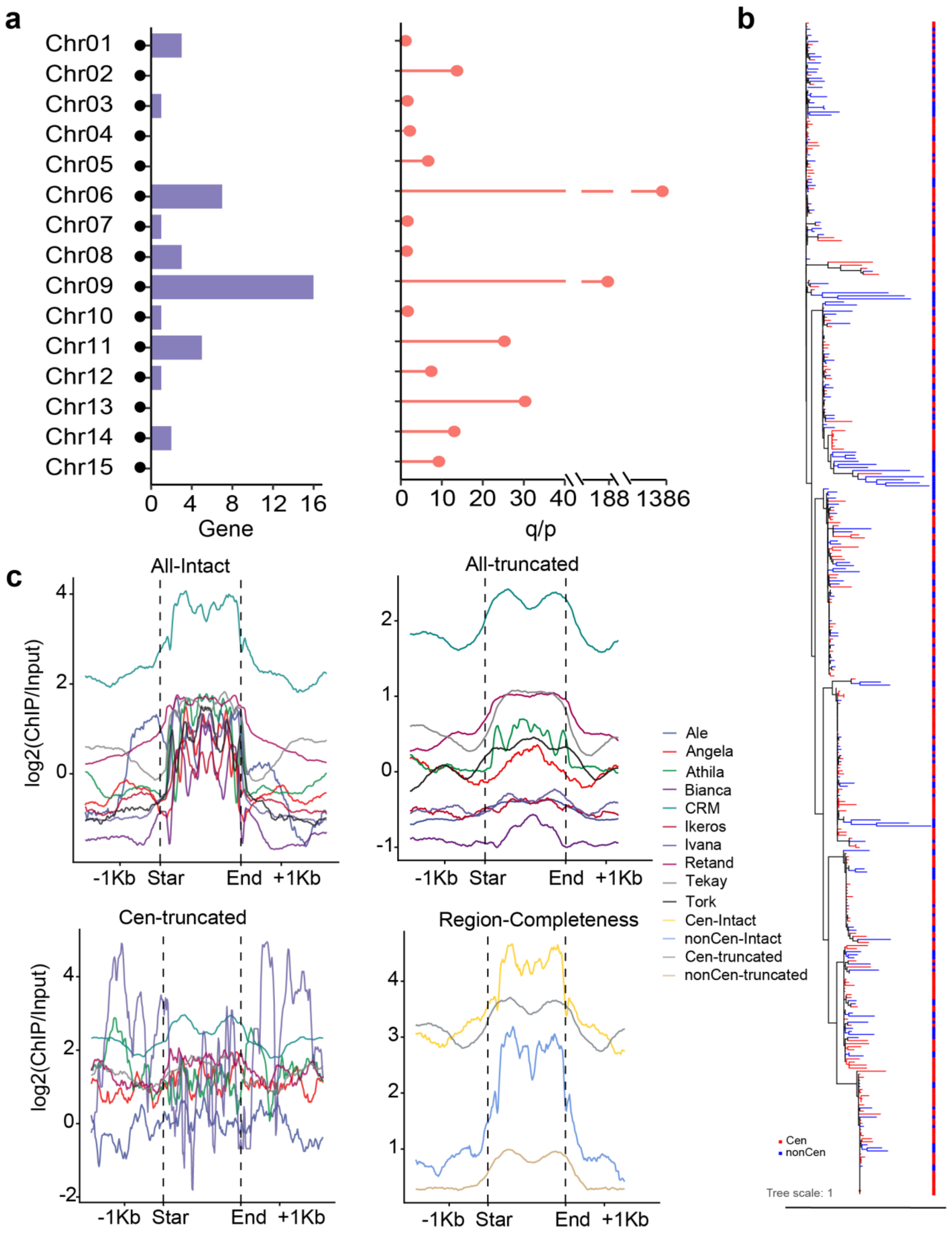


**Supplemental Figure 3. | Integrated characterization of centromere sequence composition features. a**, Number of genes at centromeres and the ratio of long arm to short arm across chromosomes. **b**, Maximum-likelihood tree based on the RT (reverse transcriptase) domains of intact CRM elements. Elements located in centromeric and non-centromeric regions are highlighted in red and blue, respectively. **c**, Comparative ChIP-seq enrichment of repeat elements in centromeric versus non-centromeric regions (Subpanels arranged left-to-right and top-to-bottom); (**i**) Genome-wide ChIP enrichment profiles of intact LTRs; (**ii**) Genome-wide ChIP enrichment profiles of truncated LTRs; (**iii**)ChIP enrichment profiles of truncated LTRs specifically in centromeric regions; (**iv**) Comparative ChIP enrichment of intact versus truncated CRMs in centromeric versus non-centromeric regions.

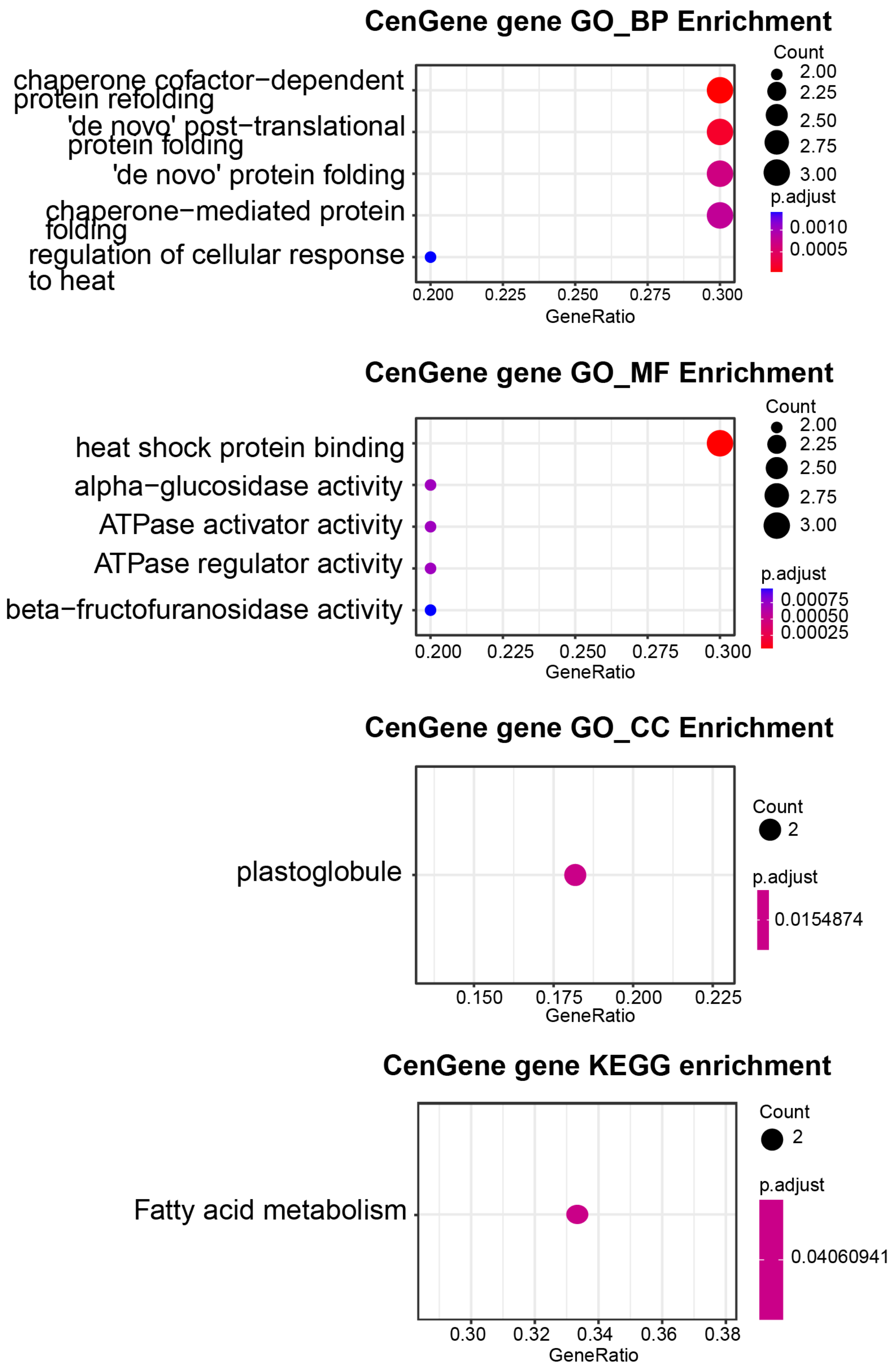


**Supplemental Figure 4. | Functional enrichment analysis of genes located at the centromere.** The circle size indicates the number of genes associated with each GO term, while color intensity reflects the p.adjust, with deeper red indicating more significance.

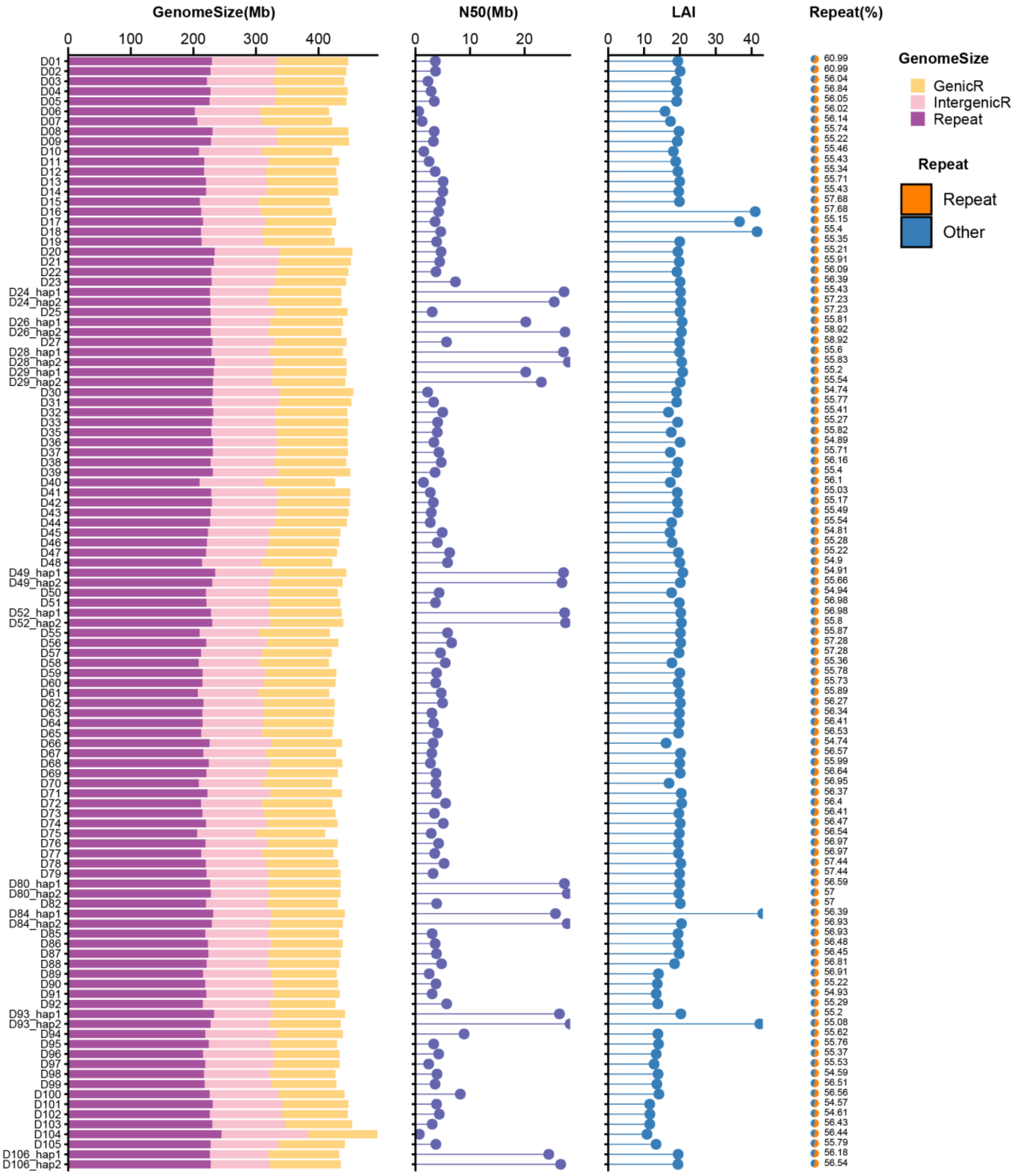


**Supplemental Figure 5. | Genome-wide features statistics.** Genome assembly, evaluation and repeat annotation of 101 accessions, including genomic length (Gb), contig N50 (Mb), LAI and repeat percentage.

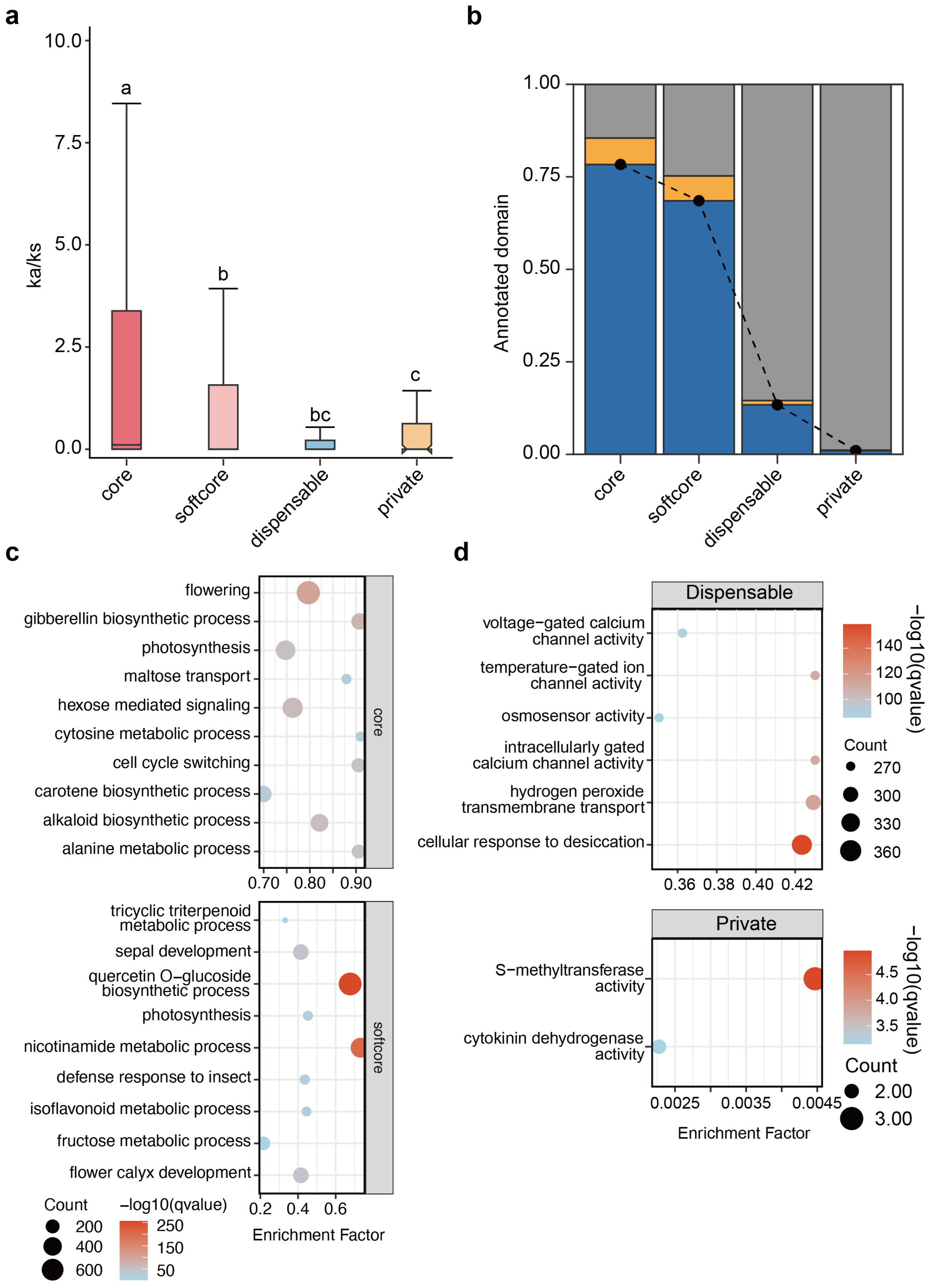


**Supplemental Figure 6. | Pan-genome analysis of 101 longan accessions. a**, Ka/Ks ratios of core, softcore, dispensable and private genes. Statistical significance was assessed using Student's t-test for multiple comparisons. **b**, Proportion of genes with Pfam domains in core, soft core, dispensable, and private genomes. Blue histograms indicate the genes with Pfam domain annotation; Orange histograms indicate the gene without Pfam domain annotation; grey histograms indicate the gene without any annotation. **c**, Gene ontology enrichment analysis of core and softcore gene families. **d**, Gene ontology enrichment analysis of dispensable and private gene families.

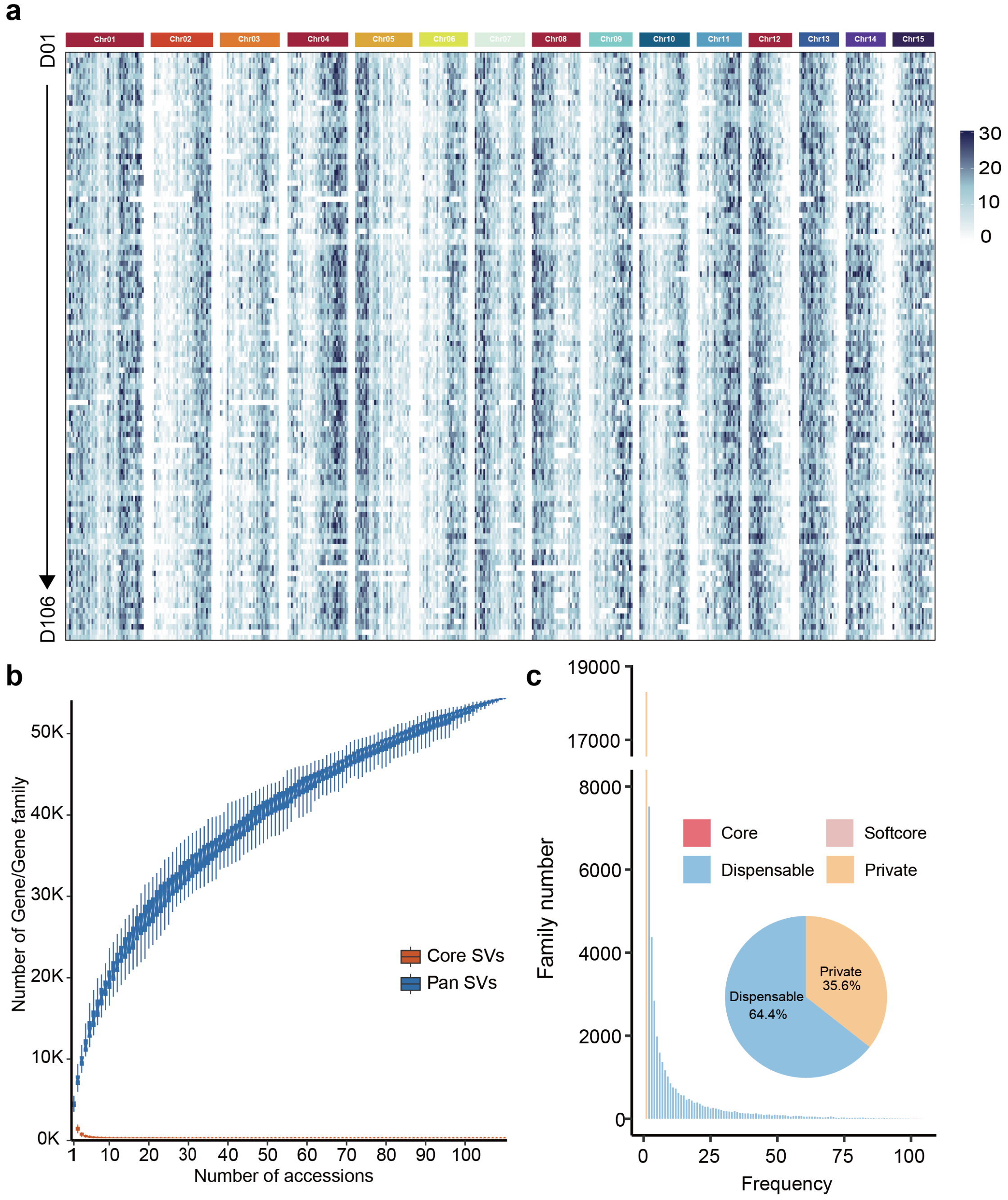


**Supplemental Figure 7. | Structural variation landscape and Pan SV analysis of longans. a**, Genome-wide density distribution of structural variants (SVs) in each individual genome. **b**, Variation of SV count in the pan- and core-SV along as additional longan genomes are incorporated. **c**, Compositions of the pan-SV and individual genomes. The histogram shows the number of SVs in the 111 genomes with different frequencies. The pie shows the proportion of the SVs marked by each composition.

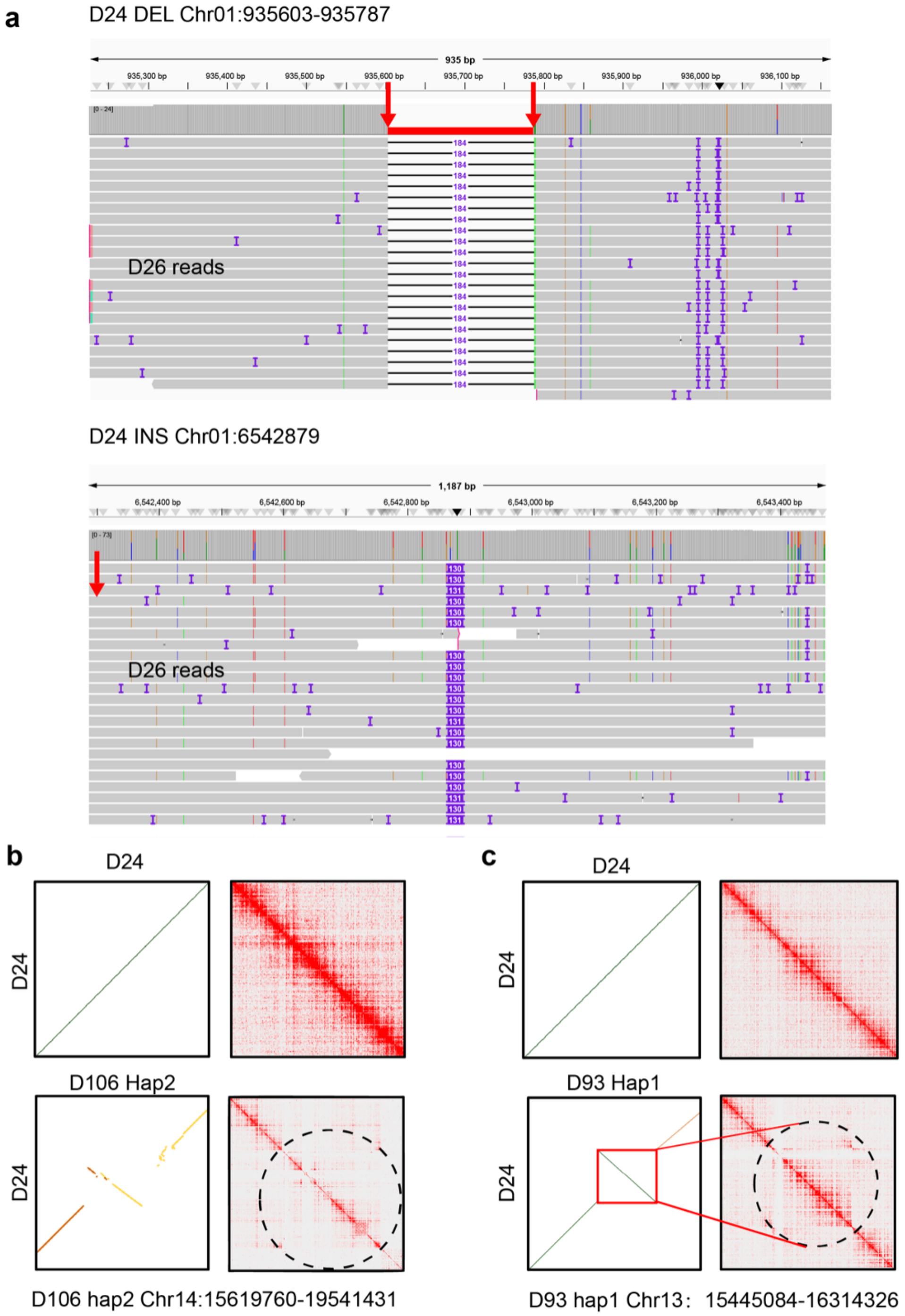


**Supplemental Figure 8. | Validation of insertion, deletion and large-scale inversion. a**, Split-read validation of representative inversion events by aligning PacBio long-reads of each accession to 'Shixia' T2T genome. The red arrows point to the breakpoints of the inversions. **b-c**, Two examples are taken as representatives. The left dot plots of genomic alignments illustrate presence of inversion, and the right Hi-C interaction matrix give the validation.

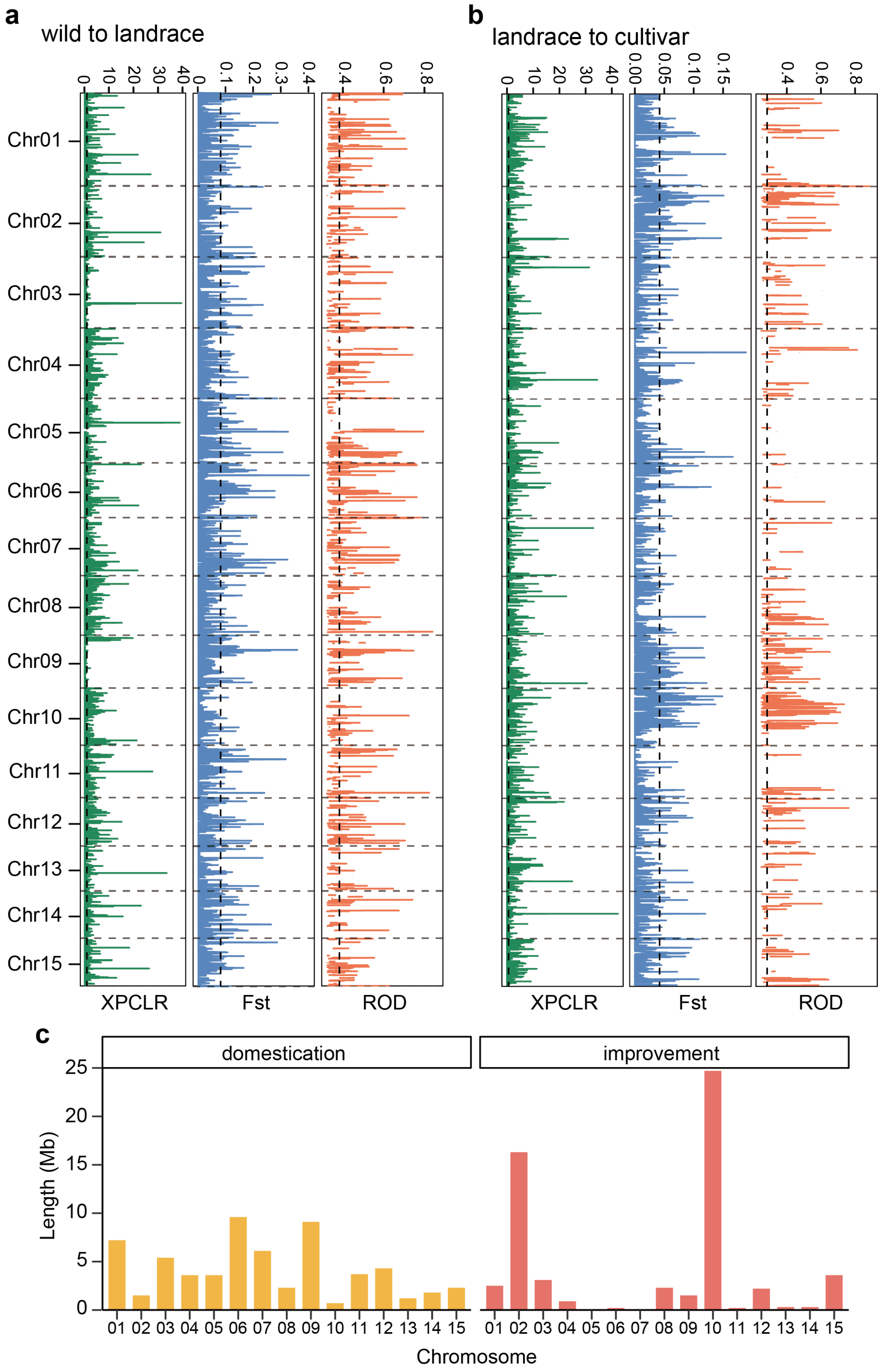


**Supplemental Figure 9. | Selective sweep in chromosomes during domestication.**

**a**, Genomic regions of selective sweep from wild accessions to landrace accessions. XPCLR, ROD, and *Fst* tests were used for selection analysis in longan. Regions of selection were selected with top 5% in at least two metrics. Vertical dashed lines indicate genome-wide threshold of top 5% selection signals (XPCLR > 0.80, ROD > 0.32 and *Fst* > 0.80). **b**, Selective sweep regions from landraces to improved accessions were detected based on the top 5% genome-wide selection signals (XPCLR > 0.66, ROD > 0.28, *Fst* > 0.42). **c**, Genomic lengths in chromosomes of selective sweep from wild accessions to landrace accessions and from landrace accessions to improved accessions.

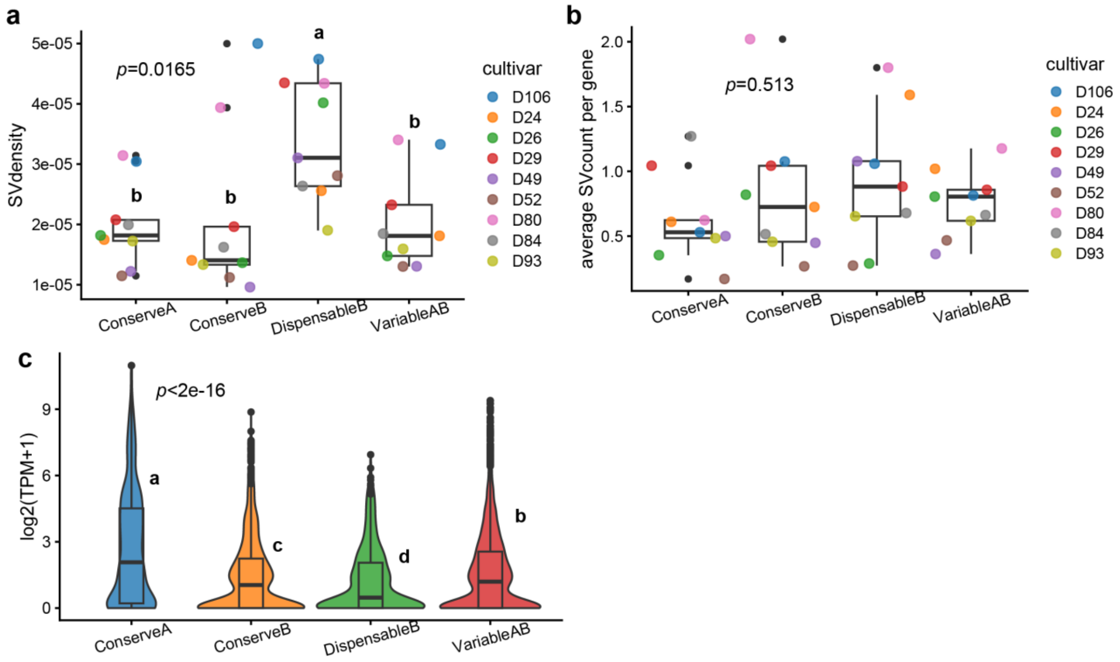


**Supplemental Figure 10.** | **SV diversity in Pan-3D. a**, SV density in different 3D structural regions. **b**, The number of SVs affecting genes in different 3D structural regions. **c**, Gene expression levels in different 3D structural regions.

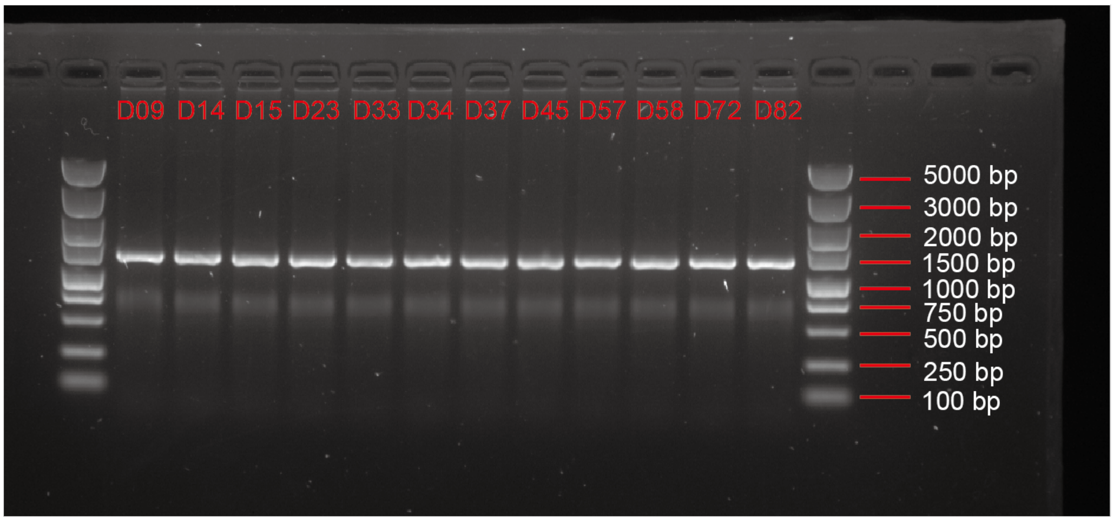


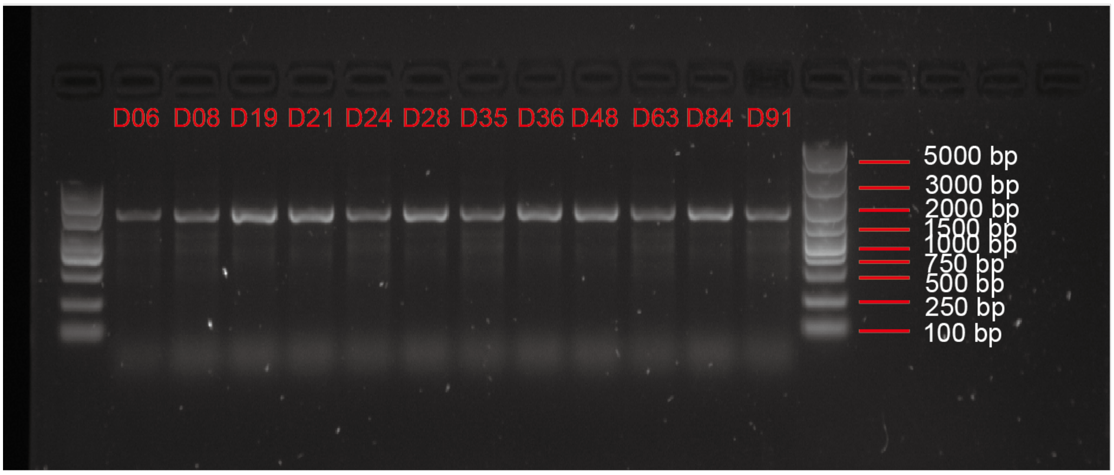


**Supplemental Figure 11. | PCR validation of detected SVs among longan accessions.** Primers were designed in 'Shixia' T2T. Genotype verification was performed at a random site, Chromosome 12: 16707284-167108690. For insertions, accessions carrying the alternative genotypes amplified larger fragments than the reference genotypes, whereas the opposite was observed for insertions.

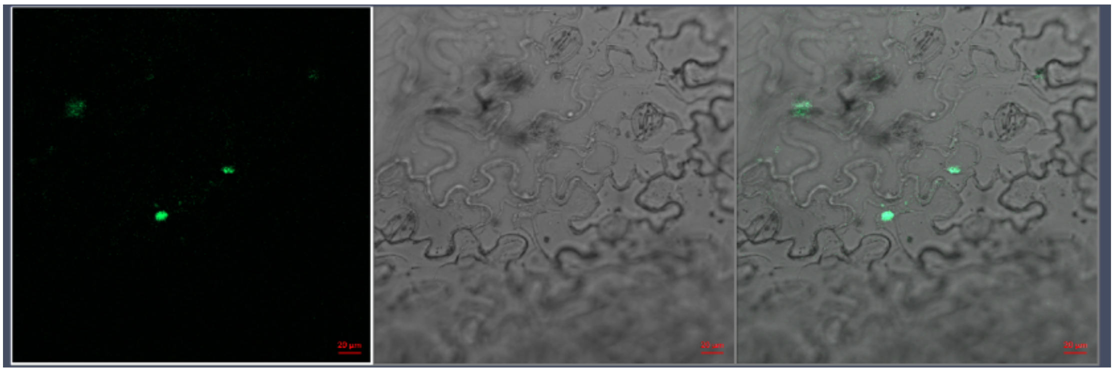

**Supplemental Figure 12. | Subcellular localization analysis of *DlDAZ* in the leaves of tobacco.**

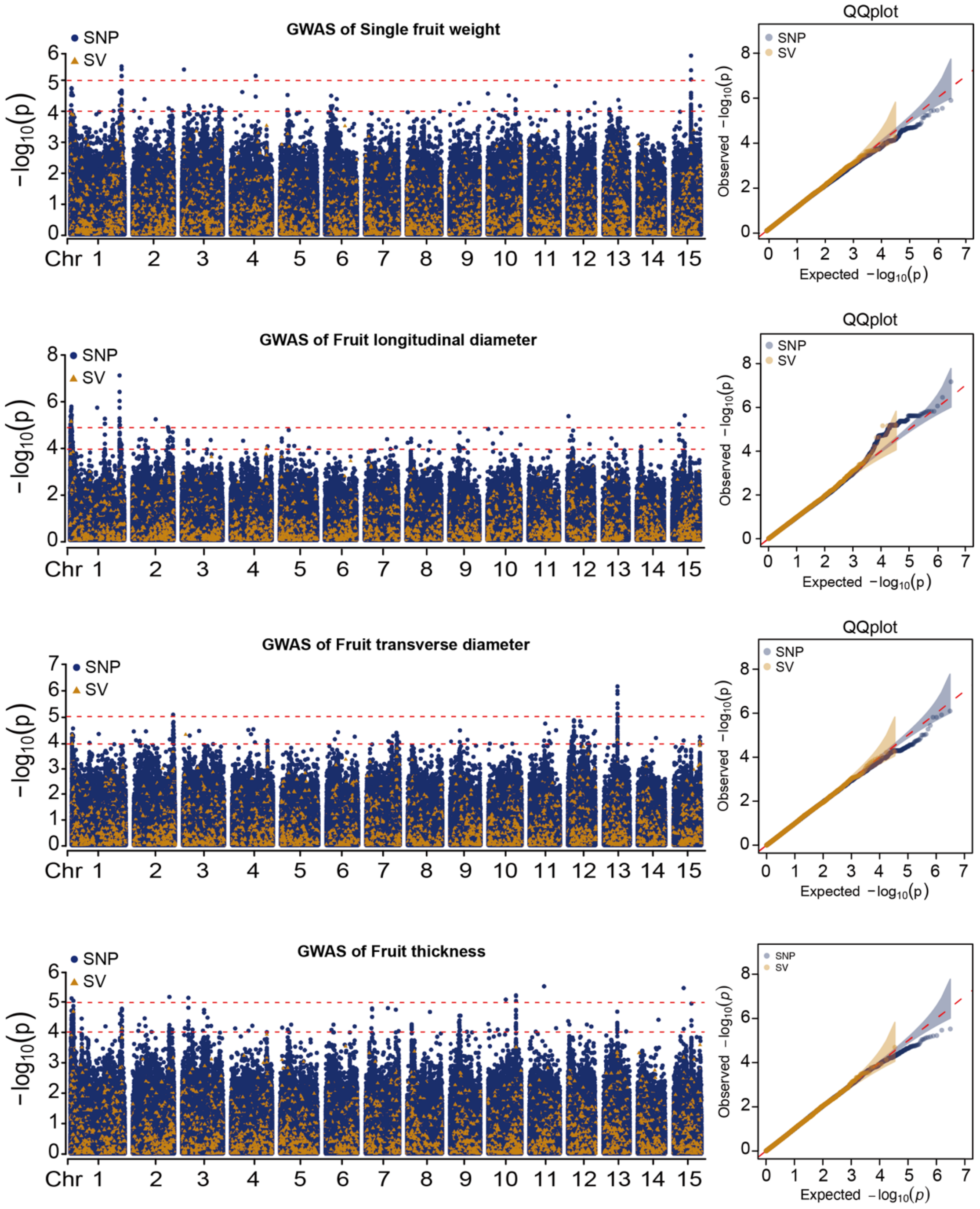


**Supplemental Figure 13. | Genome-wide association (GWAS) of fruit quality traits in longan.** Examples are given to show the SNP/SV GWAS results of single fruit weight, fruit longitudinal diameter, fruit transverse diameter and fruit thickness phenotype. Manhattan plot and corresponding QQ plot of GWAS. red dashed lines indicate SNP and SV significant threshold ($p = 1 \times 10^{-5}$ and $1 \times 10^{-4}$, respectively).
